# Migrators Within Migrators: Exploring Transposable Element Dynamics in the Monarch Butterfly, *Danaus plexippus*

**DOI:** 10.1101/2021.09.28.462135

**Authors:** Tobias Baril, Alexander Hayward

**Affiliations:** Centre for Ecology and Conservation, University of Exeter, Penryn Campus, Cornwall, TR10 9FE, UK

**Keywords:** Transposon, TE Annotation, Lepidoptera, *Danaus plexippus*, Butterfly, Genomic Deletion, Repeat, Genome Evolution

## Abstract

**Background:** Lepidoptera (butterflies and moths) are an important model system in ecology and evolution. A high-quality chromosomal genome assembly is available for the monarch butterfly (*Danaus plexippus*), but it lacks an in-depth transposable element (TE) annotation, presenting an opportunity to explore monarch TE dynamics and the impact of TEs on shaping the monarch genome.

**Results:** We find 6.21% of the monarch genome is comprised of TEs, a reduction of 6.85% compared to the original TE annotation performed on the draft genome assembly. Monarch TE content is low compared to two closely related species with available genomes, *Danaus chrysippus* (33.97% TE) and *Danaus melanippus* (11.87% TE). The biggest TE contributions to genome size in the monarch are LINEs and *Penelope*-like elements, and three newly identified families, *r2-hero_dPle* (LINE), *penelope-1_dPle* (*Penelope*-like), and *hase2-1_dPle* (SINE), collectively contribute 34.92% of total TE content. We find evidence of recent TE activity, with two novel Tc1 families rapidly expanding over recent timescales (*tc1-1_dPle*, *tc1- 2_dPle*). LINE fragments show signatures of genomic deletions indicating a high rate of TE turnover. We investigate associations between TEs and wing colouration and immune genes and identify a three-fold increase in TE content around immune genes compared to other host genes.

**Conclusions:** We provide a detailed TE annotation and analysis for the monarch genome, revealing a considerably smaller TE contribution to genome content compared to two closely related *Danaus* species with available genome assemblies. We identify highly successful novel DNA TE families rapidly expanding over recent timescales, and ongoing signatures of both TE expansion and removal highlight the dynamic nature of repeat content in the monarch genome. Our findings also suggest that insect immune genes are promising candidates for future interrogation of TE-mediated host adaptation.

## Background

Transposable elements (TEs) are autonomous DNA sequences that can move within the genome. TEs are present in nearly all eukaryotes and are important in shaping their genomes[1–8]. TEs also represent a large reservoir of sequences with the potential to contribute to host genomic novelty, and they have been implicated in the evolution of regulatory networks [3,9–13], chromosomal rearrangements [3,14–19], exon shuffling [20–22], and donation of coding sequence [20,22–27].

Due to the great diversity and dynamic nature of TEs, species-specific TE libraries and accompanying TE annotations are required to provide an understanding of their evolution and impact. By examining differences in the diversity and abundance of TEs within a genome, as well as their integration histories, proliferation frequencies, and rate of removal, we can begin to unravel the factors that govern TE evolutionary dynamics. Curating TEs in a wide range of species is also an important step more generally, to expand understanding of variation in the interplay between TEs and host genomes, and the consequences of this for host evolution [28]. Meanwhile, from a practical perspective, accurate and reliable repeat annotation is essential to provide high-quality genome annotations, and to help avoid repeat sequences being incorrectly annotated as host genes and vice versa, especially given the vast numbers of genomes currently being sequenced across the tree of life.

Lepidoptera (butterflies and moths) are an important model system in ecology and evolution. Lepidopterans have captivated researchers for decades, proving a fascinating and practical system for research into topics such as development, physiology, host-plant interactions, coevolution, phylogenetics, mimicry, and speciation (e.g. [29–33]). Over recent decades, Lepidoptera has become an important model system in genomics especially. For example, Lepidoptera genomes have been pivotal in investigations into the genomic bases of migration [34–36], warning colouration [35], adaptive radiations [33], and evolutionary studies into TEs, including TE horizontal transfer [37–39], the influence of TEs on genome size [40], and the role of a TE in the evolution of industrial melanism in the peppered moth [41].

The monarch butterfly, *Danaus plexippus*, is famous for its incredible North American migration, global dispersal, and striking orange warning colouration [35], and has been the focus of much research and broad public interest for decades. A draft monarch genome was released in 2011, and work during the last decade has focussed on revealing the genetic bases of several key traits [35, 36]. Most recently, a high-quality chromosomal assembly of the monarch genome was generated [42]. Additionally, genome assemblies for the black veined tiger (*D. melanippus*) and African Queen (*D. chrysippus*) provide a comparative context to examine monarch genomics. Yet, despite being an important research focus, we lack a detailed TE annotation for the monarch, and an accompanying understanding of monarch TE dynamics and host-TE interactions.

Here, we provide a thorough annotation of TEs within the monarch genome, alongside analyses of TE diversity and abundance, interactions with host genes, genomic distribution including hot and cold spots, and TE evolution. We find evidence of considerable recent TE activity in the monarch, with two novel Tc1 element families rapidly expanding over recent timescales (*tc1-1_dPle* and *tc1-2_dPle*). In contrast, we find that historically a pattern of high proliferation rates of LINE and *Penelope-like* elements (PLEs) was dominant, with these elements still dominating the monarch TE landscape. We also reveal the considerable role of genomic deletions in removing LINE TEs from the genome. Overall, we provide an overview of the broadscale dynamics that have shaped the TE landscape of the monarch genome, and provide a high-quality, curated, and species-specific TE annotation library for use by the scientific community.

## Results

### Transposable Element Landscape

Repeats are a major determinant of genome size [40,43–47]. However, in the monarch, we find that repeats comprise just 6.21% of the genome (total assembly size = 248.68Mb, assembly scaffold N50 = 9,210kb, 97.5% Complete Busco orthologs) (Figure 1, Table 1). This differs considerably in comparison to the TE annotation performed on the original monarch genome assembly, where repeats were estimated to cover 13.06% of the genome (Table S1; Additional File 1) [36]. When examining the source of this variation, the major difference in TE content can be attributed to unclassified elements, for which we find coverage is reduced by 16.31Mb (6.6% of total genome size) (Table S1; Additional File 1). This difference is likely due to two issues: (i) improvements in TE annotation tools over the last decade, where improved knowledge of TE structure has led to exclusion of erroneous sequences previously annotated as putatively TE, particularly given that the original annotation used similar tools and databases to those applied in this study [36]; (ii) We apply a conservative approach to TE annotation that excludes very short fragments which cannot be confidently identified as TE sequence, and we remove overlapping TE annotations.

**Figure 1.**
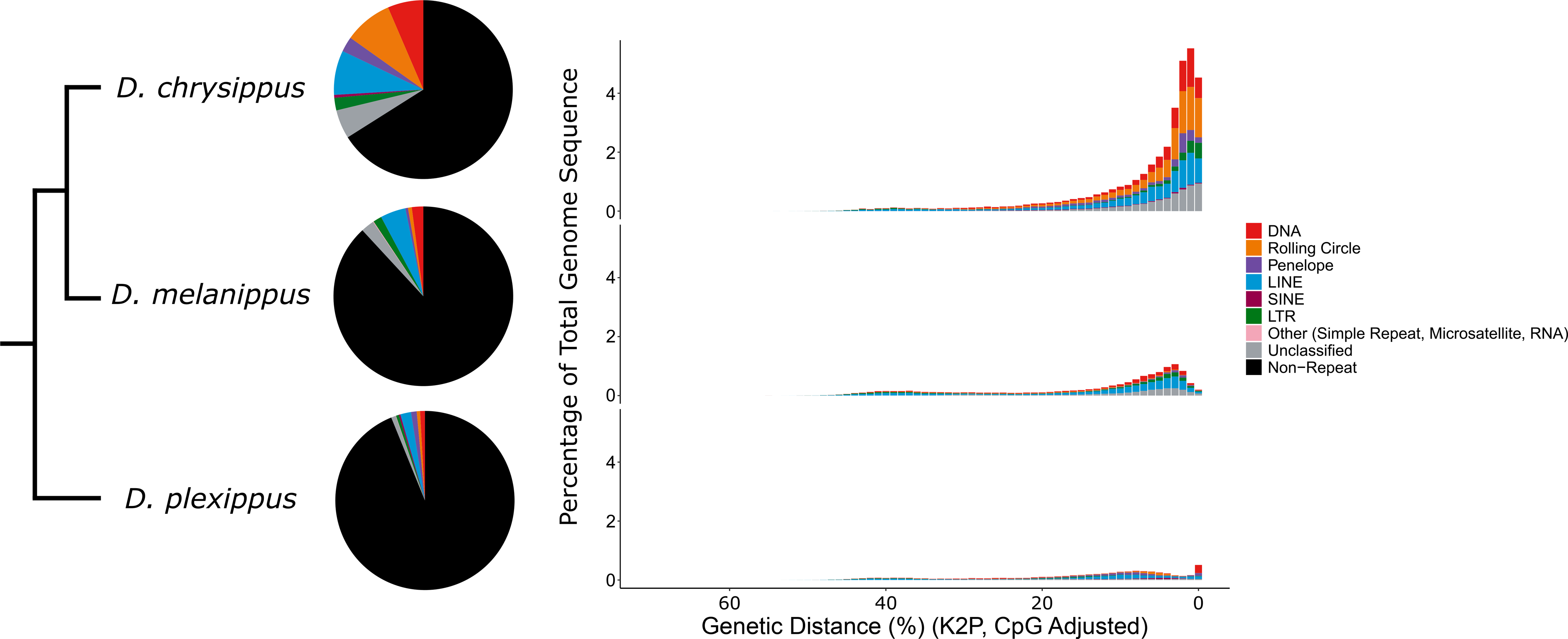
TE content in the monarch, D. chrysippus and D. melanippus. Major TE types are represented by different colours indicated in the key (A) Pie charts illustrating proportions of each Danaus genome comprised of the main TE classifications. (B) Repeat landscapes for each Danaus species. The x axis indicates the level of Kimura 2-parameter genetic distance observed between TE insertions and their respective consensus sequences in percent. More recent elements are located to the right of the x axis. The y axis indicates the percentage of the genome occupied by TE insertions of each genetic distance class.

**Table 1.**
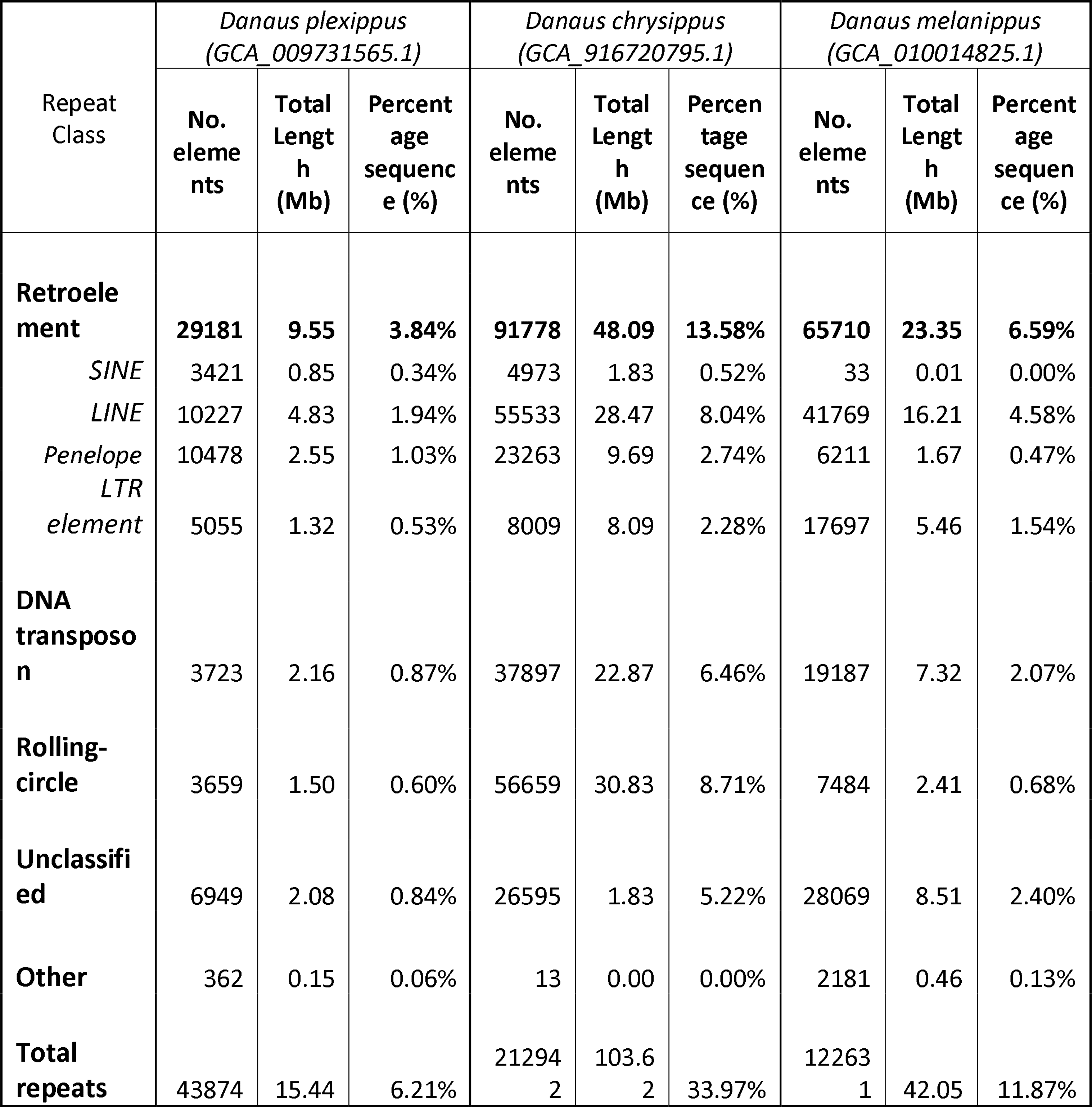
Transposable element content for the three Danaus species with available genome assemblies.

Improvements in genome assembly quality will also have improved repeat detection. Consistent with this, a standard ‘no frills’ RepeatMasker run using the Arthropoda RepBase database [48] and the Dfam database of repetitive DNA families [49] identified a repeat content of 5.60%, which is much closer to our finding of 6.21% repeat content using the earlGrey pipeline (https://github.com/TobyBaril/EarlGrey)[50], and much lower than that identified in the original TE annotation (Table S2; Additional File 1). Thus, many of the sequences annotated as unclassified in the original annotation were most likely not TE sequences and are no longer annotated as such by current annotation tools. We are, however, unable to confirm this for certain since the original *de novo* repeat library generated for the draft genome was not provided alongside the genome. We also acknowledge the considerable difference in TE content annotated in the monarch genome in this study compared to a recent study focussing on *D. chrysippus* which also annotated TEs in the monarch genome [51]. When examining the source of this variation, we find this can be attributed to the more conservative and detailed approach taken here, where for example, 95,956 extremely short annotations identified in [51] did not reach the length and score thresholds for annotation in this study, whilst we also used a monarch-specific repeat library for annotation (See Additional File 2 for full discussion).

We annotated the two additional genomes available in the genus *Danaus* using the same automated TE annotation method that we applied for the monarch. In comparison, the monarch exhibits a much lower TE content than the other two species, with repetitive elements covering 33.97% of the *D. chrysippus* genome (assembly size = 354.02Mb, contig N50 = 11.45Mb, 98.2% complete Busco orthologs), and 11.87% of the *D. melanippus* genome (assembly size = 354.18Mb, scaffold N50 = 890kb, 86.0% complete Busco orthologs) (Figure 1, Table 1). Relatively recent TE activity is indicated in all three *Danaus* species, with several TE classifications exhibiting low genetic distance to their respective consensus sequences, but there is evidence of a much greater burst of recent TE activity in. *D. chrysippus* (Figure 1). The genome size of both *D. chrysippus* and *D. melanippus* is 354Mb, whilst the monarch genome is 30% smaller at only 248Mb. Increases in TE content are apparent for all TE classifications for *D. chrysippus* and *D. melanippus* compared to the monarch, except for a decrease in PLE and SINE coverage in *D. melanippus* relative to the monarch (Table 1). The most populous TEs in *D. chrysippus* are rolling circle TEs (8.71% of genome, 30.8Mb), whilst LINEs dominate in *D. melanippus* (4.58% of genome, 16.2Mb) (Table 1).

The lower TE content observed in the monarch genome presumably arose by either: (i) loss of repeat content in the monarch; (ii) independent gains of repeat content in both *D. chrysippus* and *D. melanippus* (or in their ancestral lineages); or (iii) a gain of repeat content in the ancestral lineage leading to *D. chrysippus* and *D. melanippus* (Figure 2A). The ancestor of the monarch and *D. chrysippus* is estimated to have diverged ∼8.3Mya [52].

**Figure 2.**
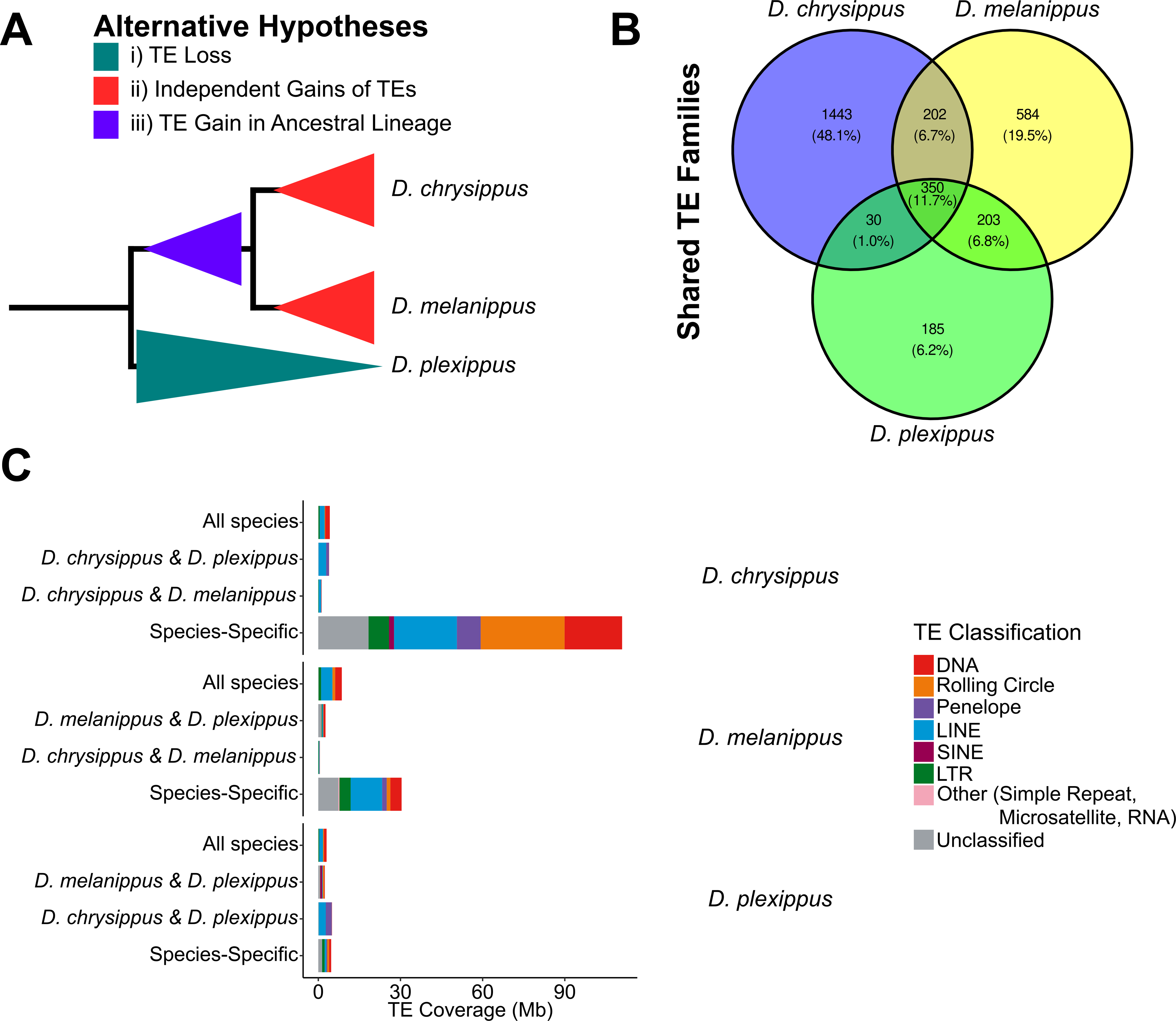
Processes leading to genome size differences between the monarch, D. chrysippus, and D. melanippus. (A) Three hypotheses leading to observed differences in genome size between Danaus species. Hypotheses indicated by colours in the key. (B) Shared and unique TE families across the three Danaus species. Numbers indicate distinct TE families. Percentages indicate proportion of total TE families found in each section of the Venn diagram. (C) TE coverage in each Danaus species split by TE shared status. Major TE types are represented by different colours indicated in the key.

Thus, using this estimate we were able to calculate DNA loss rates for the two species, to examine if the smaller genome of the monarch is a result of an increased rate of DNA loss (Table S3; Additional File 1). The estimated rates of micro- (<30bp) and mid-size (30bp to 10,000bp) deletions in the monarch and *D. chrysippus* are comparable. However, the monarch exhibits similar DNA loss rates for both micro- and mid-size deletions, whilst *D. chrysippus* has a reduced rate of DNA loss from mid-size deletions offset by an increased rate of DNA loss from micro-deletions (monarch µ=21.27 del_nt/10kb/My, micro- deletions=20.57 del_nt/10kb/My, mid-size deletions=21.96 del_nt/10kb/My; *D. chrysippus* µ=20.61 del_nt/10kb/My, micro-deletions=26.61 del_nt/10kb/My, mid-size deletions=14.62 del_nt/10kb/My). This suggests observed differences in the quantity of TE sequence between the monarch and *D. chrysippus* are due to increases in TE content in *D. chrysippus,* rather than higher rates of DNA loss in the monarch. We were unable to calculate comparative DNA loss rates for *D. melanippus*, since a divergence estimate was not available for this species. However, mean estimated DNA loss was reduced in *D. melanippus* compared to the other two species (*D. melanippus*=158.19 del_nt/10kb, monarch=176.50 del_nt/10kb, *D. chrysippus*=171.11 del_nt/10kb). *D. melanippus* exhibited a similar level of DNA loss from micro-deletions to the monarch, but a reduction compared to *D. chrysippus*, while *D. melanippus* and *D. chrysippus* exhibit a reduction in mid-size deletions compared to the monarch (Table S3; Additional File 1). Meanwhile, considerable expansions in species-specific TE families are evident in *D. chrysippus* and *D. melanippus* (Figure 2B & C). Therefore, we suggest that hypothesis (ii) is the most likely explanation for observed differences in TE load among *Danaus* species, whereby *D. chrysippus* and *D. melanippus* have both experienced independent expansions in TE content (Figure 2).

Factors that may have been responsible for these gains, particularly the large gain in TE content in the genome of *D. chrysippus*, are unclear.

Transposable Element Landscape TEs are unevenly distributed across the monarch genome, as indicated by a high standard deviation in TE base pairs per 100kb genomic window (mean = 6.1kb/100kb, SD = 4.1kb/100kb), whereby a standard deviation of 0 indicates TEs are evenly distributed across the genome (Figure 3, Figure 4, Table S4; Additional File 1). TEs were also unevenly distributed among different chromosomes (Kruskal-Wallis, χ =286.43, p<0.01). As expected, a strong positive correlation between TE coverage and chromosome size was observed (Pearson’s Correlation, r = 0.76, t_28_ = 6.2604, p < 0.01). The highest density of TEs is present on chromosome 28 (10.1kb/100kb, chromosome size = 3.99Mb), whilst chromosome 9 exhibits the lowest TE density, with less than half that observed for chromosome 28 (4.5kb/100kb, chromosome size = 9.62Mb) (Table S4; Additional File 1).

**Figure 3.**
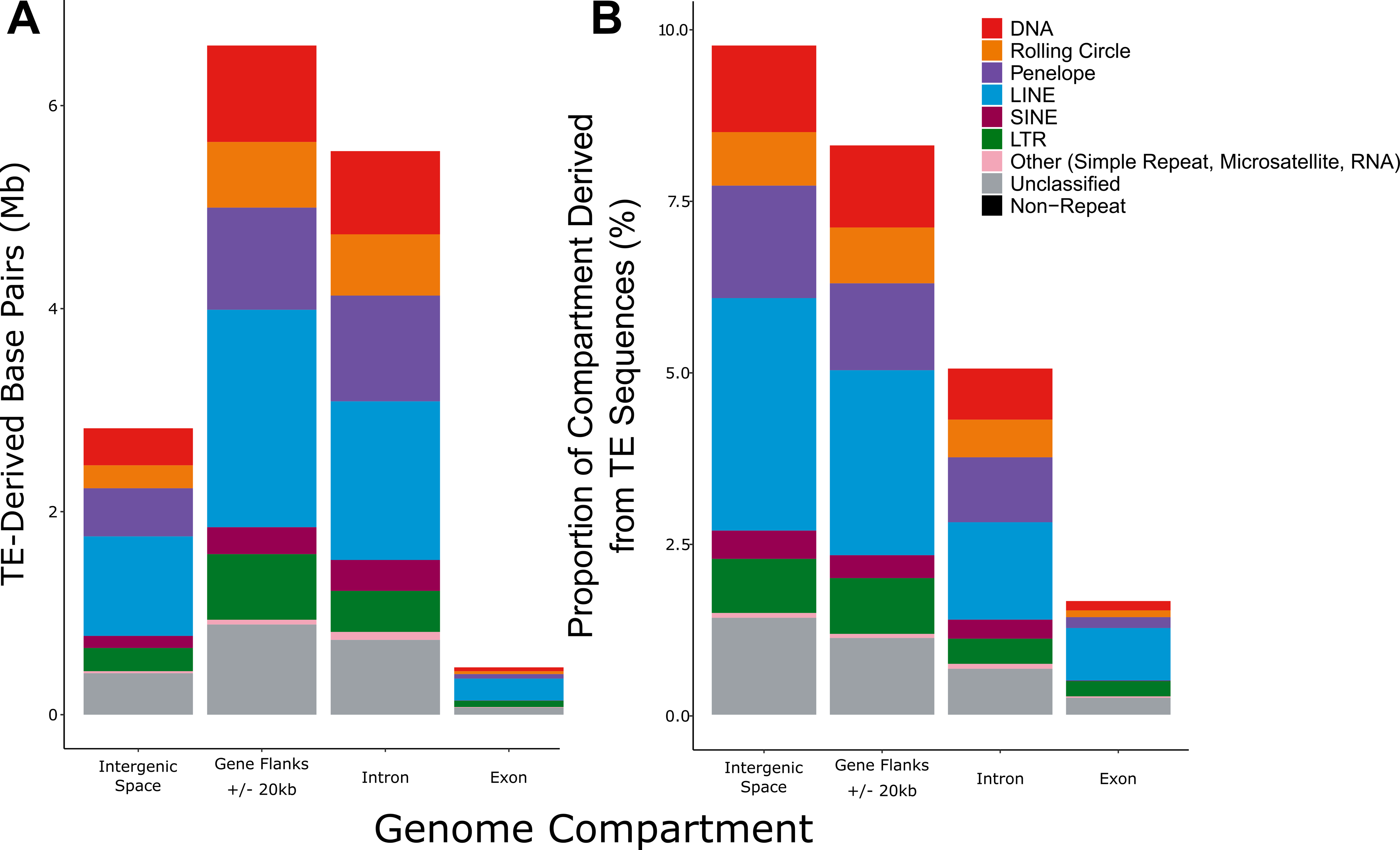
TE location in different genomic compartments in the monarch. Major TE types are represented by different colours indicated in the key. (A) Quantity of TE base pairs found in different genomic compartments. (B) Proportion of each genomic compartment contributed by TEs.

**Figure 4.**
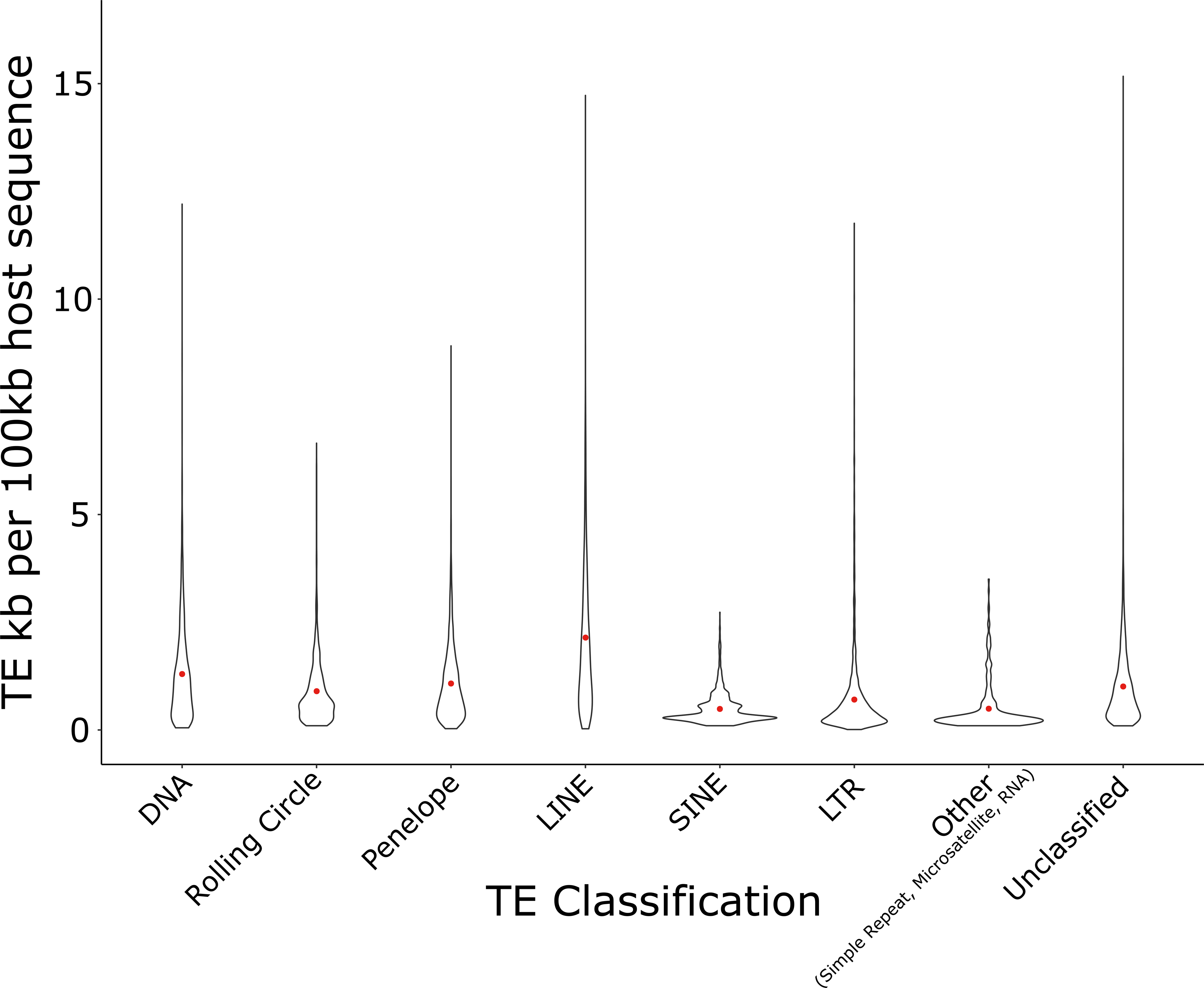
Violin plot showing kilobases of TE sequence per 100kb window across the genome for each of the main TE classifications. Red points show the mean kilobases of TE sequence per 100kb.

Similarly, TEs are unevenly distributed across the genome of *D. chrysippus* (mean = 34.0kb/100kb, SD = 28.5kb/100kb) (Table S4; Additional File 1), which also has a chromosomal-level assembly. Unlike the monarch, a weaker correlation between TE coverage and chromosome size was observed (Pearson’s Correlation, r = 0.43, t_28_ = 2.4974, p < 0.05). The highest density of TEs is found on chromosome 29 (69.8kb/100kb, chromosome size = 10.47Mb), whilst chromosome 1 exhibits the lowest TE density, with a nearly 6-fold decrease compared to chromosome 29 (12.6kb/100kb, chromosome size = 16.60Mb. In both the monarch and *D. chrysippus,* there is a significant negative correlation between chromosome size and TE density, indicating that a larger proportion of small chromosomes is composed of TE sequence (Pearson’s Correlation, monarch: r = -0.70, t_28_ = 5.1335, p < 0.01, *D. chrysippus*: r = -0.43, t_28_ = 2.5395, p < 0.05). We were unable to compare TE density in *D. melanippus* as a chromosomal-level genome assembly is not available for this species.

TEs are known to be unevenly distributed between different genomic compartments, with TE densities varying between genic and intergenic space [1]. In the monarch genome, we find that the greatest amount of TE sequence is found in gene flanking regions (6.59Mb), followed by introns (5.56MB), intergenic regions (2.82Mb) and, finally, exons (0.47Mb) (Figure 3A). While when considered as a proportion of genomic compartment size, intergenic regions have the greatest proportion of TE content (9.67%), followed by gene flanking regions (8.22%), introns (5.00%), and exons (1.64%) (Figure 3B). The relative lack of TEs in exonic regions is expected due to the potential for deleterious effects arising from transposition into coding sequence [1, 53]. LINEs and PLEs are the most populous TEs present in the monarch genome. LINEs comprise 1.94% of the monarch genome, accounting for 3.36% of intergenic sequences, 2.67% of gene flanking regions, 1.41% of intronic sequences, and 0.76% of exonic sequences (Figure 3B). Meanwhile, PLEs make up 1.03% of the monarch genome, accounting for 1.63% of intergenic sequences, 1.26% of gene flanking regions, 0.94% of intronic sequences, and 0.16% of exonic sequences (Figure 3B).

Overall, we observed a weak negative correlation between gene density and TE density in the monarch genome (Spearman’s rank, rho = -0.373, p < 0.01), suggesting that regions that contain a high density of host genes are less likely to contain TEs. We identified 25 hotspots of TE accumulation in the monarch genome, defined as 100kb windows with a TE content above a 99% area under the curve (AUC) cut-off (see Methods). This equates to windows where TE density is at least 3.03 times higher than expected (i.e., expected TE density assuming an even spread of TEs across 100kb windows = 6.15kb, lowest hotspot density = 18.64kb, highest hotspot density = 28.02kb) (Figure 5). Hotspot windows represent regions of the genome able to tolerate high densities of TE insertions. Conversely, the presence of 36 cold spots, defined as 100kb windows with TE content below a 1% AUC cut-off, where TE density is at least 12.42 times lower than expected (lowest cold spot density = 0.11kb, highest cold spot density = 0.5kb), highlights regions of the genome where very few TE insertions are tolerated. Whilst numerous TE coldspots are found on the Z chromosome, no TE hotspots are found there (Figure 5). Of note, we detected 6 separate 100kb windows labelled as TE coldspots and gene hotspots (Figure 5). Gene ontology analyses suggest that genes present in gene hotspots that are TE coldspots are involved in RNA binding, splicing, polyadenylation, mRNA stability, and translation initiation [54]; cell polarisation [55]; neuronal development; antenna and eye development [56–60]; and olfactory processes, which are hypothesised to be involved in social behaviour [36] (Table S5; Additional File 1). Given the importance of these functions, it is likely that TE insertions in these regions would be severely detrimental to host fitness, thus individuals where TEs insert into these regions may be non-viable and purged early in development.

**Figure 5.**
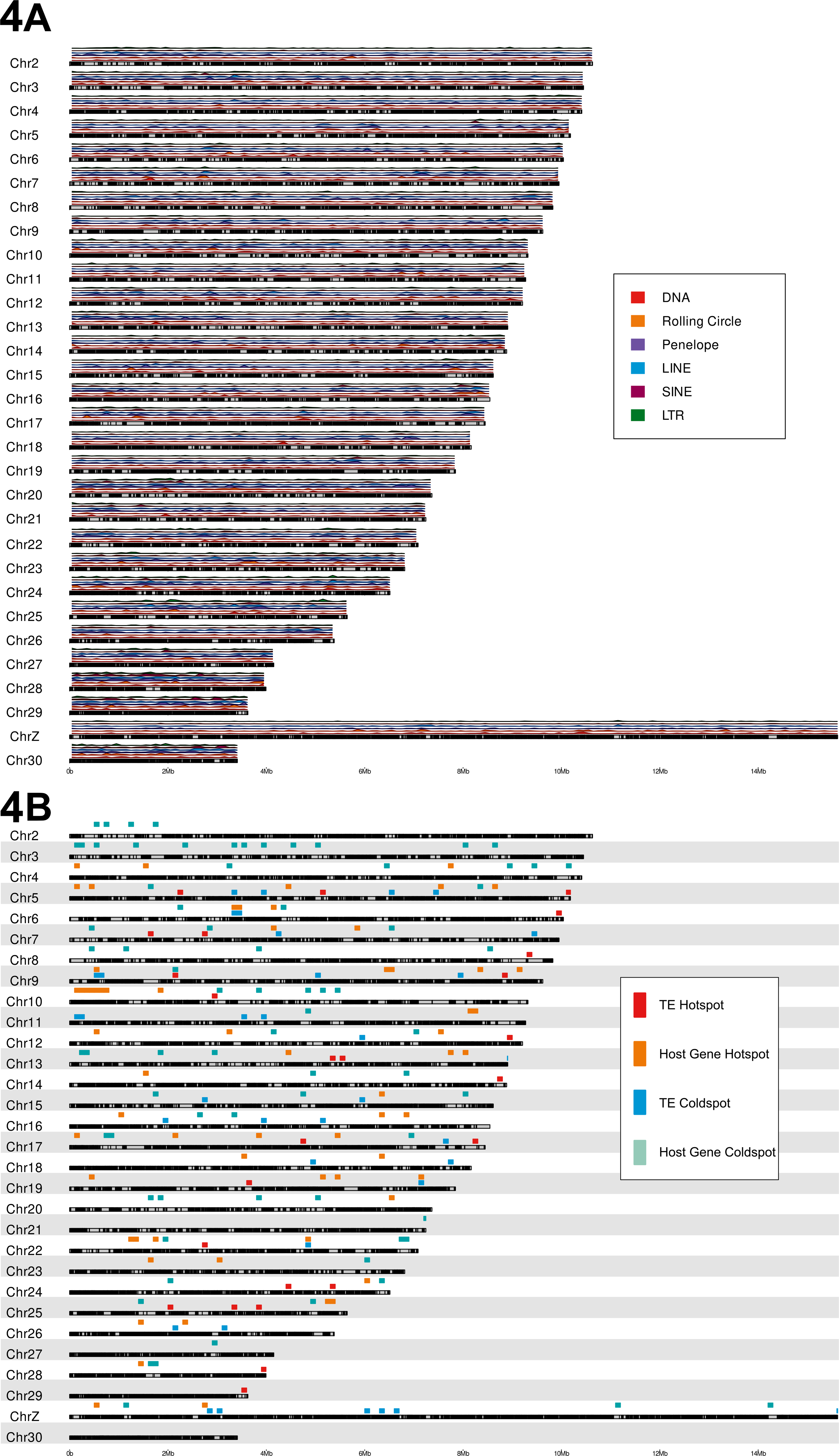
Karyoplots illustrating monarch chromosomes. Grey boxes represent chromosomes, with black regions representing genic regions of each chromosome. (A) Above each chromosome, TE density is shown for 100kb windows, with major TE types represented by different colours indicated in the key. (B) Above each chromosome, TE hotspots and coldspots are shown, with severity represented by the different colours indicated in the key,

### Transposable Element Activity

The genome of *D. plexippus* is dominated by three highly successful novel TE families, which collectively comprise 34.9% of total TE content (Table 2, Table S6; Additional File 1). Of these families, the consensus sequence of the PLE, *penelope-1_dple*, is intact and putatively transposition competent, containing both a GIY-YIG endonuclease domain and a reverse transcriptase-like (RT-like) domain, whilst the LINE element, *r2-hero-1_dple*, is putatively transposition-competent, with an intact RT-like domain (Table S7, Additional File 1). SINEs are nonautonomous elements containing an internal polymerase III promoter to facilitate their transposition [61]. As such, there are no internal coding regions to determine the transposition competency of this element, however *hase2-1_dple* is putatively full-length at 306bp.

**Table 2.** Weighted average age (TE age accounting for copy number at each divergence) and copy number of the three dominant TE families responsible for 34.9% of total TE content in the monarch.

| TE Family | TE Classification | Weighted Average Age (Mya) | Copy Number |
| --- | --- | --- | --- |
| penelope-1_dple | Penelope-like Element | 10.74 | 9901 |
| hase2-1_dple | SINE/HaSE2 | 6.39 | 2865 |
| r2-hero-1_dple | LINE/R2-Hero | 8.58 | 5432 |

Only three TE families are estimated to have inserted <1Mya in the monarch genome (Table S8, Figure S1; Additional Files 1 and 3). The low estimated age of these insertions is a sign of recent host colonisation, potentially via horizontal transposon transfer (HTT) from another host genome [1,62–65]. Peaks of low genetic distance to TE consensus sequence indicate an abundance of closely related elements and thus recent TE proliferation (Figure 1, Table S9 and Figure S2; Additional Files 1 and 4). There has been a recent burst of activity in two Tc1 elements: *tc1-1_dple* (418 copies, from ∼0.43Mya), and *tc1-2_dple* (216 copies, from ∼0.35Mya) (Table S8; Additional File 1). The rapid expansion of these Tc1 elements over such a short time period highlights their great success within the monarch genome when compared to other relatively young TEs, which are found at much lower copy numbers (Figure 6 and Figure 7, Table S8 and Table S9; Additional File 1). This recent expansion of DNA TEs is interesting given the historic dominance of LINEs in the monarch genome (1.94% of the genome, 31% of total TE content). Whilst there are more DNA TE families than LINE families in the monarch genome (319 vs 213), LINEs have been more successful in proliferating and colonising the genome than their DNA TE counterparts to date, although the reasons for this are unknown, it appears that these new DNA TE families may challenge the historic trend (Figure 6 and Figure 7).

**Figure 6.**
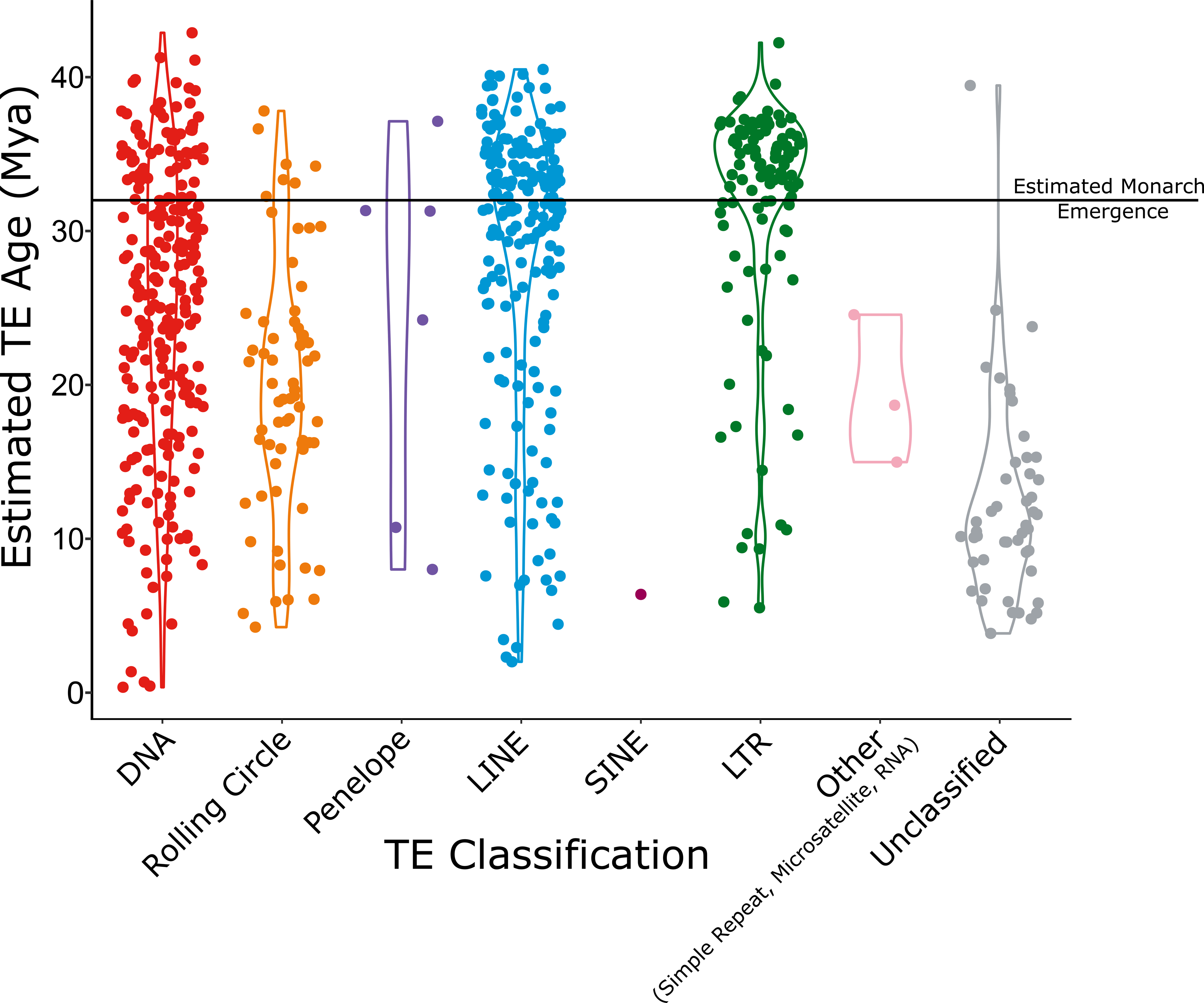
Violin plot illustrating the estimated insertion time (Mya) for each of the major TE types, represented by different colours indicated in the key. The black bar shows the estimated divergence time of the monarch, approximately 32Mya [66].

**Figure 7.**
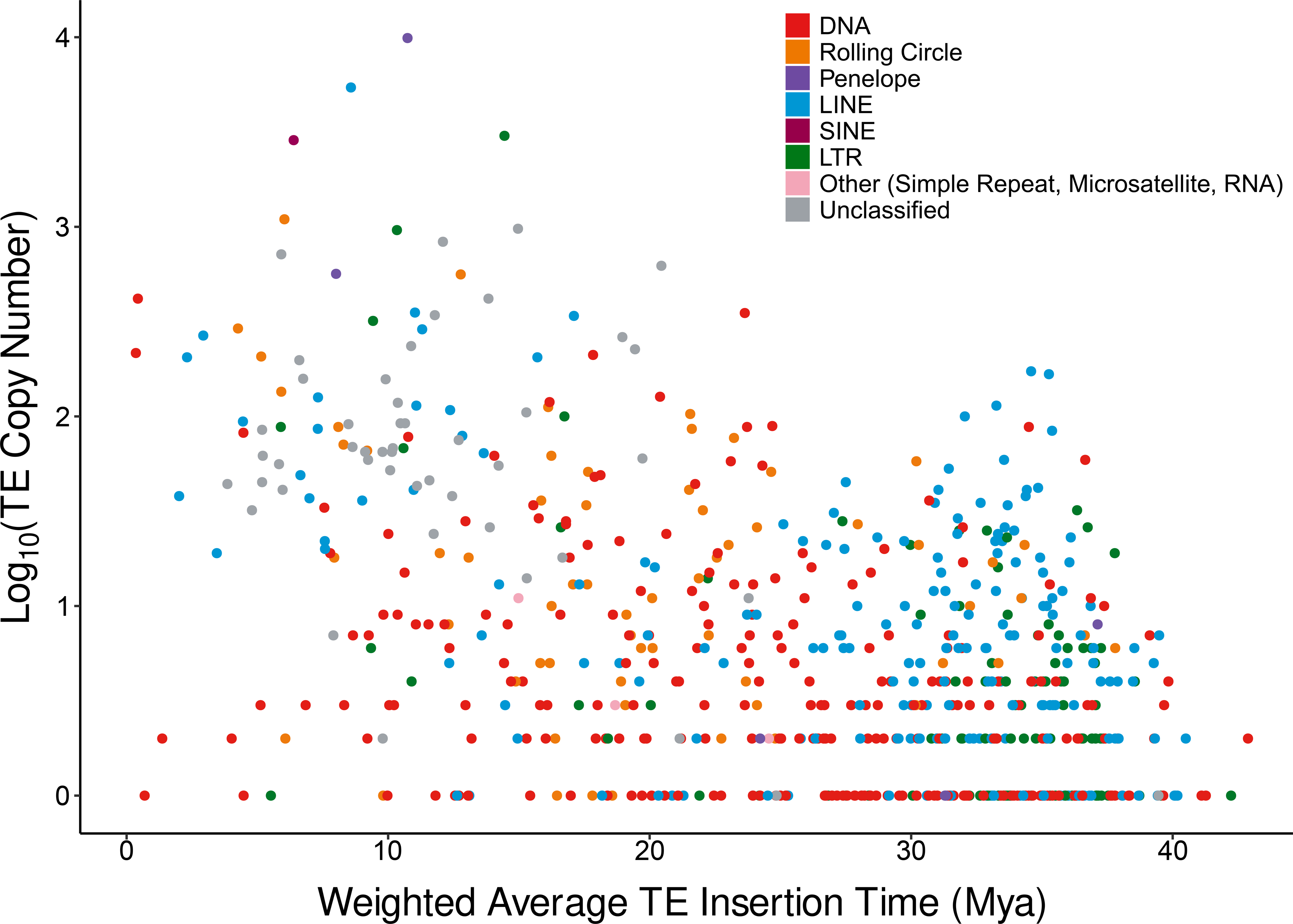
TE age against copy number. Major TE types are represented by different colours indicated in the key. X axis shows estimated age of each TE family, Y axis shows log10 transformed TE copy number identified in the monarch.

The oldest TE insertions present in the monarch genome were estimated to have occurred ∼43 Mya, predating the emergence of the monarch, which diverged ∼32 Mya [66], with the first invaders being DNA and LTR elements (Figure 6, Figure 7, Table S9, Figure S1; Additional Files 1 and 3). Thus, these TEs appear to have proliferated in an ancestral lineage after the Nymphalidae family emerged, ∼91Mya (CI: 71-112Mya) [66]. However, most TE families (79%) are estimated to have inserted after the emergence of the monarch (mean TE family age = 25.41Mya) (Figure 6, Figure 7)

### Turnover of LINEs and Penelope-like Elements

LINEs and PLEs are reverse transcribed from their 3’ end during insertion of a new copy and are often incomplete at the 5’ end due to the premature dissociation of reverse transcriptase (RT), or the activity of cellular RNases [67, 68]. In contrast, the absence of a complete 3’ region is unlikely to arise due to either process and is evidence of a genomic deletion, presumably as a consequence of ectopic recombination between homologous LINE or PLE sequences [67, 68].

Consistent with observations in the postman butterfly *Heliconius melpomene* [68] also from the family Nymphalidae, we find LINE and PLE fragments across all LINE and PLE families exhibit patterns suggestive of genomic deletions acting to remove these elements from the genome (Table 3, Figure S3; Additional File 5). In total, just under half of all LINE/PLE insertions (49.37%) are truncated at the 3’ end. For the *CR1*, *I-Jockey*, *RTE-BovB*, and *Penelope* families, most fragments display patterns suggestive of removal via genomic deletions (i.e. truncated at the 3’ end), whilst the majority of fragments of the *L2, R1*, *R2-Hero*, and *RTE-RTE* families display patterns indicative of processes associated with degradation of LINEs during mobilisation, such as premature disassociation of RT or breakdown via cellular RNAse activity (i.e. truncated at the 5’ end) (Table 3). The range of insertion times between LINEs exhibiting an abundance of 3’ fragments and those exhibiting an abundance of 5’ fragments overlap, suggesting that insertion time does not explain the inter-family differences in fragmentation (Table S9, Additional File 1). Swathes of small fragments across all LINE and PLE families (mean length = 356bp, n = 20,706) suggest a high rate of turnover of these non-LTR elements within the monarch genome, in contrast to patterns observed in mammals whereby a low rate of turnover leads to an accumulation of ageing TEs [68, 69].

**Table 3.** Number of LINEs and PLEs classified by fragment type, and putative removal process. 3’ encompasses element fragments containing a lone 3’ end, or those with 3’ ends and internal regions, and indicates premature RT dissociation or activity of cellular RNAses. 5’ encompasses element fragments containing a lone 5’ end, or those with 5’ ends and internal regions, and indicates genomic deletions.

| **LINE Classification** | **Fragment Types** | **Count** | **Proportion of Total Elements** |
| --- | --- | --- | --- |
| **CR1** | 3' | 7 | 0.13 |
|  | 5' | 48 | 0.87 |
| **I-Jockey** | 3' | 64 | 0.41 |
|  | 5' | 92 | 0.59 |
| **L2** | 3' | 138 | 0.54 |
|  | 5' | 119 | 0.46 |
| **R1** | 3' | 266 | 0.76 |
|  | 5' | 85 | 0.24 |
| **R2-Hero** | 3' | 2971 | 0.85 |
|  | 5' | 521 | 0.15 |
| **RTE-BovB** | 3' | 201 | 0.47 |
|  | 5' | 220 | 0.53 |
| **RTE-RTE** | 3' | 138 | 0.67 |
|  | 5' | 68 | 0.33 |
| **LINE (Unknown Family)** | 3' | 2 | 0.06 |
|  | 5' | 34 | 0.94 |
| **Penelope** | Lone 3', Internal + 3' | 537 | 0.15 |
|  | Lone 5', 5' + Internal, Internal Lacking 3' | 3029 | 0.85 |

Despite evidence of loss, LINE and PLE sequences still dominate the TE landscape of the monarch. To be autonomous, an intact LINE must contain start and stop codons and an RT domain [70], whilst an intact PLE must contain an RT domain and a GIY-YIG endonuclease domain [71]. In total, we identified five intact LINEs and two intact PLEs in the monarch genome (Table S7, Additional File 1), including one copy of *r2-Hero-1_dPle* and 2 copies of *penelope-1_dPle*, which are the two most populous TE families in the monarch genome, suggesting continued activity (Table 2). We identified 4 intact LINEs in *D. melanippus* and 9 in *D. chrysippus*, however in both species intact PLEs were absent. We also observed accumulations of certain LINE families in the monarch, however only 11.38% of LINEs and 3.51% of PLEs have maintained a length above 1kb, falling short of the expected length given an intact LINE ORF encoding RT is ∼3kb [72] (Table S10; Additional File 1). Short elements are more likely to persist in the genome than their longer counterparts, as they are less prone to recombination [19]. This, combined with the high activity level of LINEs and PLEs before and throughout the evolution of the monarch (Figure 3), explains the dominance of LINE and PLE fragments across the genomic landscape, despite their high rates of turnover [19, 68].

### Association of TEs with Monarch Colouration and Immune Genes

Given a previous role for TEs in wing colouration in the peppered moth (*Biston betularia*) [41] and the clouded yellow butterfly (*Colias croceus*) [73], we investigated the possibility of TE associations with wing colouration in the monarch. The myosin gene is associated with wing colouration in the monarch [35], and we identified a single unclassified element 7kb upstream of this gene (Figure S4; Additional File 6). This location contrasts to that of TEs implicated in colouration in the peppered moth and clouded yellow, where the TE insertions were found within an intron, and 6kb downstream of the gene body, respectively [41, 73]. When compared to all monarch genes, the association of a single TE with the myosin gene is within the expected range to find within 20kb of a genic locus (See Methods), and therefore is not evidence of TE enrichment (Figure S5, Additional File 7).

Immune genes are among the fastest evolving metazoan genes given strong selection arising from exposure to harmful pathogens and parasites [74–77], and so are key candidates for exploring the role of TEs in host evolution. Across the monarch genome, TEs of all major types were found within immune gene flanks (20kb either side of gene bodies). Regions surrounding immune genes contain a three-fold increase in TE content when compared to genes involved in other functions, although this level of enrichment is just above a 5% level of significance (Welch’s T-Test, T_9.0915_ = 2.1351, p = 0.061). However, a significant difference in TE abundance between different types of immune genes was apparent (Kruskal-Wallis ^2^_38_ = 104.88, p < 0.01), suggesting an enrichment for TEs around Class II-associated invariant chain peptide (CLIP)-like modulation genes when compared to immune genes involved in IKK gamma, Tab2, SOCS, and IMD signalling. (Figure S6; Additional File 8).

When comparing associations between major TE types around immune genes, we find that LINEs and PLEs are significantly more abundant than rolling circles (Kruskal-Wallis, χ _7_ = 29.88, p < 0.01), but there were no significant pairwise differences between the remaining major TE types. However, LINEs and PLEs are not enriched around immune genes compared to genome-wide levels (LINEs: genome-wide = 1.94% of genome size, immune gene regions = 2.04% of immune gene compartment size; PLEs: genome-wide = 1.03% of genome size, immune gene regions = 0.96% of immune gene compartment size).

## Discussion

We provide a detailed annotation of TEs in the monarch genome, an analysis of their diversity and evolution, an investigation of their influence in shaping the monarch genome, and a comparative overview of TE load for the monarch and two congeneric species. We find a large reduction in TE sequence compared to the original annotation performed on the draft monarch genome. We suggest that this is due to many sequences in the original annotation no longer being recognised as TEs when using updated TE databases and current annotation tools. However, we are unable to confirm this, due to the original *de novo* TE library and annotation results being unavailable. This raises an important issue in genomics currently. Given TE annotation is a key step in genome assembly and annotation, we suggest that it should be required to include TE libraries alongside the publication of genome projects, ideally in a specific repository, to: (i) reduce duplication of efforts required for reannotation; (ii) improve reproducibility in annotation; and (iii) improve TE consensus libraries to reduce cases of a single TE family being given different names in multiple species. As the number of sequenced genomes increases, the wider availability of TE libraries will benefit a wide range of studies, where gene annotations can be improved by accurate TE annotation. The availability of large numbers of well-curated TE libraries will also facilitate large-scale studies of TE evolutionary dynamics, and the impacts of TEs on specific traits of interest, and host evolution more generally.

We identify an uneven distribution of TEs across the genome. TE insertions in non-coding regions are less likely to have a detrimental impact on host genome function than TE insertions into coding regions [1,21,53,78–80], and so are less stringently removed by selection, enabling TE accumulation in non-coding regions. Indeed, some TEs actively avoid coding regions by targeting regions of the genome unlikely to impact host fitness [53, 81], with this process potentially occurring in the monarch, where we find overlapping gene hotspots and TE cold spots. TE abundance is highest in intergenic regions. Of note, the highest TE coverage is found in host gene flanking regions rather than intergenic regions distal from host genes, where the risk of detrimental impacts of TE insertion are thought to be reduced [53, 81]. The finding is interesting given the potential involvement of TEs in the evolution of gene regulatory networks [10–13,82] and the uneven distribution of TEs across the monarch genome. Although this could also be due to the relatively compact nature of the monarch genome, where genic regions account for 56% of the genome and host gene flanks account for a further 32%, leaving only 12% of the genome distal from host genes (i.e intergenic). TEs are implicated in rapid increases in intron size in eukaryotic genomes, where short non-autonomous DNA TEs independently generate hundreds to thousands of introns [1,53,83]. It is also hypothesised that TE-derived introns are responsible for the prevalence of nucleosome-sized exons (DNA sequences ∼150bp that can wrap around a single histone core) observed in eukaryotes [83]. Thus, the presence of TEs in intronic sequences in the monarch genome could reflect the status of introns as refugia where TE insertions are not as likely to be purged, or alternatively it could indicate ongoing intron generation, with TE insertions leading to the generation of new, or longer, introns over evolutionary time.

The presence of several young high copy number TE families in the monarch genome demonstrates the success of TEs in naïve host genomes [81,84–87]. The putative recent arrival of two novel Tc1 families (*tc1-1_dple* and *tc1-2_dple*), has led to a wave of TE proliferation in the monarch genome. These elements are closely related and located in Clade 130 of Tc1/mariner phylogeny, which contains elements from diverse insect species, including butterflies, aphids and ants, in a recent phylogenetic study of DDE TE diversity [88] (Figure S7; Additional File 9). Ants and aphids are known to share environments with monarch larvae on milkweed plants [89], raising the potential for HTT events between these groups [64, 90]. Elucidating the origin of recent Tc1 insertions in the monarch genome will require a fuller comparison across related butterfly species and other host plants and insects that are known to share environments with the monarch, as more genomes become available. Over time, it is hypothesised that the activity of the novel families identified will decrease as the host develops mechanisms to suppress activity, either through the capture of TE sequences in piRNA regions, or through other defensive mechanisms such as methylation [87,91,92]. A reduction in transposition of DDE transposase-containing DNA transposons can also be attributed to increases in non-autonomous copies [88]. In the meantime, however, these TEs are proliferating and dynamically altering the genomic landscape of *D. plexippus*, with potential consequences for ongoing genome structure and functional evolution [1–3,19].

Factors leading to the relatively low TE content observed in the monarch compared to other *Danaus* species are currently unknown. The maintenance of a small genome in the monarch may be partly attributed to the high turnover of TEs, evidenced by a lack of ancient peaks in repeat landscapes and the presence of few intact LINEs and PLEs, many of which show hallmarks of genomic deletions. We did not observe large differences in the number of intact LINEs between the three *Danaus* species considered here, however both *D. chrysippus* and *D. melanippus* lack transposition competent PLEs. In *D. melanippus*, there are only small numbers of intact LINEs and a relative lack of recent LINE activity, despite historical LINE activity having made the largest TE contribution to genome size (Figure 1 and Figure 2). In contrast, there is evidence of very recent LINE activity in *D. chrysippus*, although much very recent TE activity is attributed to DNA and rolling circle elements. (Figure 1 and Figure 2).

Ectopic recombination between similar elements is likely to play a significant role in the removal of LINEs and PLEs. The only LINE or PLE families with an abundance of 3’ fragments are: *L2*, *R1*, *R2-Hero*, and *RTE-RTE*. While it is tempting to suggest that L2, *R1*, *R2-Hero*, and *RTE-RTE* insertions may have a lower impact on host fitness than other LINE insertions, and so are “allowed” to degrade over time, rather than being removed *en masse* through genomic mechanisms, it should be noted that ectopic recombination is a passive process. However, removal via ectopic recombination does not account for inter-family differences in the ratio of fragments undergoing degradation through ectopic recombination versus degradation via the activity of RT or cellular RNAses. In the case of the most abundant family, *R2-Hero*, these elements may have so far avoided ectopic recombination, given these elements are relatively young compared to others in the same classification (insertion ∼8.58Mya, mean LINE insertion time 28.51Mya ) (Figure 6 and Figure 7, Table S9; Additional File 1)). Our results are consistent with those of previous findings in *Heliconius* species [93], in which a high TE turnover and a relative lack of more ancient TEs were observed, suggesting these characteristics might be widespread among nymphalids or lepidopterans more generally, although further comparative studies will be required to confirm this. In addition to TE turnover, TE-driven increases in genome size are likely also limited through inhibition of TE activity via epigenetic mechanisms and piRNAs [1,91,94,95]. Further research to characterise piRNA clusters in the monarch, and interrogation of its transcriptome, will further improve our understanding of how these processes have shaped its TE landscape.

The lack of TE hotspots on the Z chromosome is perhaps unexpected given sex chromosomes are thought to accumulate TEs faster than autosomes due to suppressed homologous recombination [96–98]. If this was the case in the monarch, one might expect to observe TEs accumulating due to the inability of the host genome to purge insertions. The Z chromosome of the monarch is the product of a fusion between an ancestral-Z and neo-Z genome segment, with distinct modes of dosage compensation acting on either section [42]. The unique chromatin remodelling dynamics on the Z chromosome in the monarch might moderate TE access to regions of the Z chromosome, although further research is required to understand the interplay between host processes and TE dynamics on this unique sex chromosome. Meanwhile, the three-fold increase in TE abundance around immune genes is intriguing and warrants future research to further understand the impact of TEs in these regions. The association with CLIP modulation genes is particularly interesting, as CLIP peptides block class II MHC binding grooves to modulate the immune response by influencing antigen presentation to CD4 T cells by class II MHC molecules [99]. Class II MHC genes are diverse and evolve rapidly, maintained by strong positive selection and frequent gene conversion [100, 101]. Given the strong selection these loci are under, these are ideal candidates for further study to investigate a potential role for TEs in the adaptive immune system. The use of TE knockout studies in combination with transcriptomic analyses may elucidate the potential impacts TEs have on immune gene evolution in the monarch.

## Conclusions

In-depth studies into TE landscapes and dynamics within host species of broad scientific interest remain relatively few, even with widespread acknowledgement of the important contributions that TEs have made to eukaryotic evolution [1,43,102]. We provide a detailed annotation of TEs in the monarch genome and analysis of TE diversity and evolution, including the identification of highly successful young DNA TE families, which mirror the activity profile of the most successful LINEs present in the genome. We provide a comparative context with two other *Danaus* species, demonstrating the considerable differences in TE content that can occur even among congeneric species. We present evidence of ongoing TE expansion and removal, highlighting the dynamic nature of repeat content within genomes over time. We also identify an increase in TE abundance around genes of immune function, providing potential candidates for future studies to elucidate the impact of TEs on immune function in the monarch. Further large-scale comparative studies will be greatly beneficial to further our understanding of the evolutionary dynamics of TEs, and the drivers leading to diverse TE landscapes across lineages.

## Materials and Methods

### Transposable Element Annotation

The monarch chromosomal assembly, Dplex_v4, was downloaded from NCBI (Refseq GCF_009731565.1), along with corresponding genome annotations in GFF format [42, 103]. The monarch genome assembly is 248.676Mb, with a GC content of 32.1%, and comprises 4,115 scaffolds with a scaffold N50 of 9.21Mb.

The *D. plexippus* genome was annotated using Earl Grey (github.com/TobyBaril/EarlGrey/) [104]. Known repeats were first identified and masked using RepeatMasker (version 4.1.0) [105] using the *Arthropoda* library from RepBase (version 23.08) and Dfam (release 3.3) [48, 49]. Low complexity repeats and RNA were not masked (*-nolow and -norna*) and a sensitive search was performed (*-s*). Following this, a *de novo* repeat library was constructed using RepeatModeler (version 1.0.11), with RECON (version 1.08) and RepeatScout (version 1.0.5) [106–108]. To generate maximum-length consensus sequences for the *de novo* repeat library, novel repeats identified by RepeatModeler were curated using an automated version of the ‘BLAST, Extract, Extend’ process [28]. A BLASTn search was performed on the *D. plexippus* genome to obtain up to the top 40 hits for each TE family identified by RepeatModeler [109]. Sequences were extracted with 1,000 base pairs of flanking sequences at the 5’ and 3’ ends. Each set of family sequences were aligned using MAFFT (version 7.453) [110]. Alignments were trimmed with trimAl (version 1.4.rev22) to retain high-quality positions in the alignment (*-gt* 0.6 *-cons* 60) [111]. Updated consensus sequences were computed with EMBOSS cons (*- plurality* 3) to generate a new TE library featuring consensus sequences with extended flanks [112]. This process was repeated through 5 iterations to obtain maximum-length consensus sequences.

Following automated curation, *de novo* family sequences were subject to a manual curation following protocols described by Goubert et al. (2022)[113]. Here, the final alignments generated by Earl Grey were visualised using AliView [114], and poorly-aligned positions and ends were trimmed where single-copy DNA was present. Consensus sequences were generated using EMBOSS cons [112] to generate a library of manually curated elements, which was clustered using cd-hit-est [115] with the options recommended by Goubert et al. (2022) [113] to reduce redundancy. To classify TEs, the TE library was analysed with the diagnostic tool TE-Aid (https://github.com/clemgoub/TE-Aid) to identify TE-associated protein domains in sequence ORFs and to look for the presence of TIRs and LTRs [105,109,112]. Following this, blastn and nhmmer searches were performed against the Dfam database (version 3.3) to identify homology to previously described elements [49,109,116].

The resulting *de novo* repeat library was combined with the RepBase *Arthropoda* library and utilised to annotate repetitive elements using RepeatMasker with a score threshold at the relatively conservative level of 400 (*-cutoff* 400), to exclude poor matches unlikely to be true TE sequences. Following this, the *D. plexippus* genome was analysed with LTR_Finder (version 1.07), using the LTR_Finder parallel wrapper [117, 118] to identify full-length LTR elements. Final TE annotations were defragmented and refined using the loose merge (*-loose*) command in RepeatCraft [119]. Overlapping annotations were removed using MGKit (version 0.4.1) filter-gff, with the longest element of overlapping annotations retained (*-c* length *-a* length) [120]. Finally, all repeats less than 100bp in length were removed before the final TE quantification to decrease spurious hits. Final repeat annotation is provided in datasets S1 and S2 (Additional Files 10 and 11), and *de novo* TE library is provided in dataset S3 (Additional File 12).

### Quantifying DNA Loss Rates Between the Monarch and *D. chrysippus*

To estimate DNA loss rates between the monarch and *D. chrysippus*, the DelGet.pl script (https://github.com/4ureliek/DelGet) was used to estimate DNA loss from small (<30nt) and mid-size (30<nt<10,000) deletions, as described previously [45]. Briefly, the script identifies orthologous regions of 10,000bp between three input species (an outgroup and two species with known branch length in Mya) using BLAT (https://genome.ucsc.edu/FAQ/FAQblat.html). These regions are extracted and aligned with MUSCLE [121] before deletions are quantified. Species-specific gaps are identified and quantified. Deletion rates are then calculated between the two species of interest by dividing the normalised deleted nucleotide count (del_nt/10kb) by the corresponding Mya since divergence.

### Identifying Transposable Element Association with Genomic Compartments

The *D. plexippus* gene annotation file was processed to generate a GFF file containing coordinates of exons, introns, and 5’ and 3’ flanking regions. Gene flanking regions are defined here as 20kb directly upstream or downstream of the gene body. These flanking regions are determined to identify TEs at a distance that could be described as the proximate promoter region, rather than just accounting for the core promoter region, to include both promoter and more distal enhancer regions for genes [122]. Bedtools intersect (version 2.27.1) was used to identify the number of bases of overlap (*-wao*) between TEs and different genomic compartments [123]. Following this, quantification was performed in R.

### TE Hotspots and Coldspots

To identify TE and gene hotspots and coldspots, chromosomal scaffolds were split into 100kb windows, and the TE density in each window was calculated and used to produce the frequency distribution of observations. A polynomial model was fitted to the frequency distribution, to generate a smooth curve through the data points using the lm function in R. Using these models, the total area under the curve (AUC) was calculated using the area_under_curve function of the bayestestR package [124]. From this, the 1% and 99% AUC cutoff values were determined. Starting with the first data point and iteratively adding subsequent datapoints with each round, the AUC for each collection of points was determined. The maximum AUC value not exceeding 99% of the total AUC was taken as the cutoff where windows containing higher TE or gene coverage were considered to be hotspots. The minimum AUC value not exceeding 1% of the total AUC was taken as the cutoff where windows containing lower TE or gene coverage were considered to be coldspots (Figure S8; Additional File 13). Karyoplots of the *D. plexippus* genome, illustrating gene and TE locations, highlighting TE hotspots and coldspots were generated using the KaryoploteR package in R [125].

### Estimating TE Age and Activity

TE activity dates were estimated using Kimura 2-parameter genetic distances (including CpG sites) calculated between individual insertions and the consensus sequence of each TE family [126]. A neutral mutation rate of 0.0116 substitutions per site per million years was used for *D. plexippus*, as previously described for the monarch [127].

### Turnover of LINEs and PLEs

To interrogate LINE and PLE decay patterns, each LINE and PLE sequence was extracted from the genome using bedtools getfasta [123]. Extracted sequences were used to query the repeat library using BLASTn with tabular output specified (*-outfmt* 6). In the event of multiple matches, only the top match was retained (defined by percent identity).

Coordinates of each match were scaled to normalise the length of each consensus sequence. Mapped fragments were plotted using ggplot2 of the tidyverse package in R [128, 129].

### Identification of Intact LINEs

To identify full-length LINEs, all elements longer than 2,700bp were extracted from the monarch genome using bedtools getFasta [123]. Open reading frames (ORFs) were detected using EMBOSS getORF [112]. ORF sequences were filtered to retain all sequences > 600aa. RPSBLAST+ (v2.9.0+) [109] was used to query the ORF sequences against the CDD database [130] to identify conserved protein domains. Sequences were classified as intact LINEs based on the presence of relevant protein domains as previously described [72] (Table S7, Additional File 1).

### Identification of Intact PLEs

To identify intact PLEs, all elements longer than 2,400bp were extracted from the monarch genome using bedtools getfasta [123] as full-length PLEs are ∼2,500bp long [131]. Open reading frames (ORFs) were detected using EMBOSS getORF [112]. ORF sequences were filtered to retain all sequences >200aa. RPSBLAST+ (v2.9.0+) [109] was used to query the ORF sequences against the CDD database [130] to identify conserved protein domains. Sequences were classified as intact PLEs based on the presence of reverse transcriptase (RT) and a GIY-YIG endonuclease.

### Classification of Novel DNA TEs

To further classify novel DNA elements in the monarch beyond homology to known elements, as used by RepeatModeler, DDE transposases were compared with curated alignments for all DDE TE superfamilies [88]. Specifically, dashes in curated alignments of all known DDE TE sequences for each of the 19 families were removed, and the resultant fasta file was used as the subject in a blastx search against the *de novo* monarch TE library, with an e-value threshold of 1x10^-20^ and tabular output specified (*-evalue* 1e-20 *-outfmt* 6) [109]. The best match for each *de novo* query sequence was retained, with a minimum 50% identity over at least 50% of the total length of the curated DDE subject sequence. Following this, novel DNA elements were only detected for the Tc1-Mariner family (Figure S7; Additional File 9). Transposase sequences for the *de novo* monarch sequences were extracted and aligned to the original DDE alignments for their respective families using the profile alignment command in MAFFT [110]. Sequences from related transposon superfamilies were used as outgroups, as described [88]. To estimate phylogenies for the two DDE superfamilies, we performed 1,000 ultra-fast bootstrap replicates in IQ-TREE (v1.6.12) [132] specifying the best-fit amino acid model for each family as indicated by ModelFinder [133]. Phylogenetic trees were visualised in figtree (v1.4.4) (http://tree.bio.ed.ac.uk/software/figtree/).

### Association of TEs with Monarch Colouration and Immune Genes

We obtained the transcript sequence in fasta format for the myosin gene responsible for wing colouration from MonarchBase, under accession DPOGS206617 [34]. To identify the matching region in the v4 monarch assembly, a blastn search was performed to query the monarch assembly [109]. The coordinates of the myosin gene were obtained and used in a bedtools window search to identify TEs within 20kb (*-w* 20000)[123]. TE copy number within 20kb of gene bodies was quantified for each monarch gene and a polynomial model was fitted to the frequency distribution to generate a smooth curve through the data points using the lm function in R. Using these models, the total area under the curve (AUC) was calculated using the area_under_curve function of the bayestestR package [124]. From this, the 2.5% and 97.5% AUC cutoff values were determined, equivalent to selecting the central 95% of the data distribution. Starting with the first data point and iteratively adding subsequent datapoints with each round, the AUC for each collection of points was determined. The maximum AUC value not exceeding 97.5% of the total AUC was taken as the cutoff where genes containing higher TE copy number were considered significantly enriched for TEs. The minimum AUC value not exceeding 2.5% of the total AUC was taken as the cutoff where genes containing lower TE copy number were considered significantly depleted for TEs (Figure S5; Additional File 7). A karyoplot of the myosin gene region including 20kb flanks was generated in R using the karyoploteR package [125].

To investigate associations between monarch immunity genes and TEs, nucleotide and protein sequences of monarch immune genes were obtained in fasta format from MonarchBase [34]. To identify corresponding regions in the v4 monarch assembly, blastn and tblastn searches were used to query the monarch genome assembly with the monarch immune gene nucleotide and protein sequences [109]. Matches were filtered by percentage identity and query match coordinates to identify the locus of each immune gene. A bed file was generated, containing the name and location of each immune gene, and a bedtools window search was performed to identify TEs within 20kb of each immune gene(*-w* 20000)[123]. The same method was applied to identify TEs within 20kb of all other genes. TE bases surrounding each immune gene were quantified in R [134]. As TE count was not normally distributed, a Kruskal-Wallis test was used to compare TE abundance between immune genes and other genes, between different TE classifications around immune genes, and between different immune gene types. Unless otherwise stated, all plots were generated using the ggplot2 package of the tidyverse suite in R [128, 129].

## Additional Files

Additional File 1: Supplementary Tables, including contents page with table legends. Tables S1-S10.

Additional File 2: Detailed discussion on the discrepancies between TE annotations of the monarch presented in this study, and those presented in another recent study focussing on *chrysippus* [51].

Additional File 3: Figure S1. Scatter plot illustrating estimated TE age for each TE family. Major TE types are represented by different colours indicated in the key. Elements are ordered from oldest to youngest, within each major type.

Additional File 4: Figure S2. Repeat landscapes for the monarch. The x axis indicates the level of Kimura 2-parameter genetic distance observed between TE insertions and their respective consensus sequences in percent. More recent elements are located to the right of the x axis. The y axis indicates the percentage of the genome occupied by TE insertions of each genetic distance class.

Additional File 5: Figure S3. Plot illustrating LINE and Penelope fragments annotated in the monarch genome. The x axis indicates normalised start and end coordinates for each element relative to its consensus. Elements are organised from longest to shortest, with short elements found towards the top of the plot. Families are represented by different colours indicated by the key.

Additional File 6: Figure S4. Karyoplot of the monarch gene association with colouration, myosin, obtained from MonarchBase under accession DPOGS206617. Main TE types within 20kb are represented by different colours indicated in the key.

Additional File 7: Figure S5. Plot illustrating the distribution of TE copy number found with 20kb of all genes, showing the 2.5% and 97.5% AUC cutoffs used to determine the TE copy number within 20kb of a given gene for it to be significantly depleted or enriched in comparison to other genes.

Additional File 8: Figure S6. Boxplots illustrating: (A) Normalised TE coverage in 20kb regions surrounding each immune gene type. (B) Normalised TE count in 20kb regions surrounding each immune gene type. Main TE types are represented by different colours indicated in the key.

Additional File 9: Figure S7. Phylogenetic tree of Tc1-Mariner DDE transposases, with novel monarch families added (highlighted in red). Branch support from 1000 ultra-fast bootstrap repetitions. Novel Tc1-Mariner families highlighted in teal are the young TEs found with extremely high copy number.

Additional File 10: Dataset S1. Coordinates and classification of all transposable element sequences identified in the monarch genome using the *de novo* library in conjunction with RepBase *Arthropoda* library and Dfam, in bed format.

Additional File 11: Dataset S2. Coordinates and classification of all transposable element sequences identified in the monarch genome using the *de novo* library in conjunction with RepBase *Arthropoda* library and Dfam, in GFF format.

Additional File 12: Dataset S3. *De novo* transposable element consensus sequences identified in the monarch in FASTA format.

Additional File 13: Figure S8. Plots illustrating: (A) Distribution of genome windows with TE coverage, with lines showing 1% and 99% AUC cutoffs used to define hotspots and coldspots. (B) Distribution of genome windows with host gene coverage, with lines showing 1% and 99% AUC cutoffs used to define hotspots and coldspots.

## Abbreviations

TE: Transposable element
Non-LTR: Non-long terminal repeat
LTR: Long terminal repeat
LINEs: Long INterspersed Elements
SINEs: Short INterspersed Elements
PLEs: *Penelope*-like Elements
TSS: Transcription Start Site

## Declarations

Ethics approval and consent to participate: Not applicable

## Consent for Publication

Not applicable

## Availability of Data and Materials

All data generated or analysed during this study are included in this article and its supplementary information files.

## Authors’ Contributions

TB developed the pipelines and scripts for automated TE annotation and downstream analyses, produced the figures, and drafted the manuscript. AH conceived and coordinated the study and participated in writing the manuscript. All authors read and approved the final manuscript.

## Acknowledgements

We would like to thank Dr James Galbraith, Ryan Biscocho, and Ryan Imrie for constructive comments on early drafts of this manuscript, and Dr Aurélie Kapusta for support in running the DNA deletions analysis.

## References

1. Bourque G, Burns KH, Gehring M, Gorbunova V, Seluanov A, Hammell M, et al. Ten things you should know about transposable elements. Genome Biol 2018;19:199. https://doi.org/10.1186/s13059-018-1577-z.

2. Klein SJ, O’Neill RJ. Transposable elements: genome innovation, chromosome diversity, and centromere conflict. Chromosom Res 2018;26:5–23. https://doi.org/10.1007/s10577-017-9569-5.

3. Feschotte C, Pritham EJ. DNA Transposons and the Evolution of Eukaryotic Genomes. Annu Rev Genet 2007;41:331–68. https://doi.org/10.1146/annurev.genet.40.110405.090448.

4. Belyayev A. Bursts of transposable elements as an evolutionary driving force. J Evol Biol 2014;27:2573–84. https://doi.org/10.1111/jeb.12513.

5. Schrader L, Schmitz J. The impact of transposable elements in adaptive evolution. Mol Ecol 2018. https://doi.org/10.1111/mec.14794.

6. Casola C, Lawing AM, Betrán E, Feschotte C. PIF-like transposons are common in Drosophila and have been repeatedly domesticated to generate new host genes. Mol Biol Evol 2007;24:1872–88. https://doi.org/10.1093/molbev/msm116.

7. Jangam D, Feschotte C, Betrán E. Transposable Element Domestication As an Adaptation to Evolutionary Conflicts. Trends Genet 2017;33:817–31. https://doi.org/10.1016/j.tig.2017.07.011.

8. Volff JN. Turning junk into gold: Domestication of transposable elements and the creation of new genes in eukaryotes. BioEssays 2006. https://doi.org/10.1002/bies.20452.

9. Mita P, Boeke JD. How retrotransposons shape genome regulation. Curr Opin Genet Dev 2016;37:90–100. https://doi.org/10.1016/j.gde.2016.01.001.

10. Bourque G. Transposable elements in gene regulation and in the evolution of vertebrate genomes. Curr Opin Genet Dev 2009;19:607–12. https://doi.org/10.1016/j.gde.2009.10.013.

11. Feschotte C. Transposable elements and the evolution of regulatory networks. Nat Rev Genet 2008;9:397–405. https://doi.org/10.1038/nrg2337.

12. Rebollo R, Romanish MT, Mager DL. Transposable Elements: An Abundant and Natural Source of Regulatory Sequences for Host Genes. Annu Rev Genet 2012;46:21–42. https://doi.org/10.1146/annurev-genet-110711-155621.

13. Chuong EB, Elde NC, Feschotte C. Regulatory activities of transposable elements: From conflicts to benefits. Nat Rev Genet 2017;18:71–86. https://doi.org/10.1038/nrg.2016.139.

14. Bennetzen JL. Transposable element contributions to plant genome evolution. Plant Mol Biol 2000;42:251–69. https://doi.org/10.1023/A:1006344508454.

15. Casals F, Cáceres M, Ruiz A. The Foldback-like transposon Galileo is involved in the generation of two different natural chromosomal inversions of Drosophila buzzatii. Mol Biol Evol 2003;20:674–85. https://doi.org/10.1093/molbev/msg070.

16. Campos-Sánchez R, Cremona MA, Pini A, Chiaromonte F, Makova KD. Integration and Fixation Preferences of Human and Mouse Endogenous Retroviruses Uncovered with Functional Data Analysis. PLoS Comput Biol 2016;12:1–41. https://doi.org/10.1371/journal.pcbi.1004956.

17. Oliver KR, Greene WK. Transposable elements: Powerful facilitators of evolution. BioEssays 2009;31:703–14. https://doi.org/10.1002/bies.200800219.

18. Hughes JF, Coffin JM. Human endogenous retroviral elements as indicators of ectopic recombination events in the primate genome. Genetics 2005;171:1183–94. https://doi.org/10.1534/genetics.105.043976.

19. Kent T V., Uzunović J, Wright SI. Coevolution between transposable elements and recombination. Philos Trans R Soc B Biol Sci 2017;372. https://doi.org/10.1098/rstb.2016.0458.

20. Coates BS, Hellmich RL, Grant DM, Abel CA. Mobilizing the genome of lepidoptera through novel sequence gains and end creation by non-autonomous Lep1 Helitrons. DNA Res 2012;19:11–21. https://doi.org/10.1093/dnares/dsr038.

21. Biémont C, Vieira C. What transposable elements tell us about genome organization and evolution: The case of Drosophila. Cytogenet Genome Res 2005;110:25–34. https://doi.org/10.1159/000084935.

22. Morgante M, Brunner S, Pea G, Fengler K, Zuccolo A, Rafalski A. Gene duplication and exon shuffling by helitron-like transposons generate intraspecies diversity in maize. Nat Genet 2005;37:997–1002. https://doi.org/10.1038/ng1615.

23. Philippsen GS, Avaca-Crusca JS, Araujo APU, DeMarco R. Distribution patterns and impact of transposable elements in genes of green algae. Gene 2016;594:151–9. https://doi.org/10.1016/j.gene.2016.09.012.

24. Wu M, Li L, Sun Z. Transposable element fragments in protein-coding regions and their contributions to human functional proteins. Gene 2007;401:165–71. https://doi.org/10.1016/j.gene.2007.07.012.

25. Daniel C, Behm M, Öhman M. The role of Alu elements in the cis-regulation of RNA processing. Cell Mol Life Sci 2015;72:4063–76. https://doi.org/10.1007/s00018-015-1990-3.

26. Vorechovsky I. Transposable elements in disease-associated cryptic exons. Hum Genet 2010;127:135–54. https://doi.org/10.1007/s00439-009-0752-4.

27. Bejerano G, Lowe CB, Ahituv N, King B, Siepel A, Salama SR, et al. A distal enhancer and an ultraconserved exon are derived from a novel retroposon. Nature 2006;441:87–90. https://doi.org/10.1038/nature04696.

28. Platt RN, Blanco-Berdugo L, Ray DA. Accurate transposable element annotation is vital when analyzing new genome assemblies. Genome Biol Evol 2016;8:403–10. https://doi.org/10.1093/gbe/evw009.

29. Bates HW. Contributions to an Insect Fauna of the Amazon Valley.-Lepidoptera:- Heliconinæ. J Proc Linn Soc London Zool 1862;6:73–7. https://doi.org/10.1111/J.1096-3642.1862.TB00932.X.

30. Müller F. Ituna and Thyridia: a remarkable case of mimicry in butterflies. Trans Entomol Soc L 1879;1879:20–9.

31. Ehrlich PR, Raven PH. Butterflies and Plants: A Study in Coevolution. Evolution (N Y) 1964;18:586. https://doi.org/10.2307/2406212.

32. Kettlewell, B. Evolution of melanism: the study of a recurring necessity 1973.

33. Dasmahapatra KK, Walters JR, Briscoe AD, Davey JW, Whibley A, Nadeau NJ, et al. Butterfly genome reveals promiscuous exchange of mimicry adaptations among species. Nature 2012;487:94–8. https://doi.org/10.1038/nature11041.

34. Zhan S, Reppert SM. MonarchBase: the monarch butterfly genome database. Nucleic Acids Res 2013;41:D758–63. https://doi.org/10.1093/NAR/GKS1057.

35. Zhan S, Zhang W, Niitepõld K, Hsu J, Haeger JF, Zalucki MP, et al. The genetics of monarch butterfly migration and warning colouration. Nat 2014 5147522 2014;514:317–21. https://doi.org/10.1038/nature13812.

36. Zhan S, Merlin C, Boore JL, Reppert SM. The monarch butterfly genome yields insights into long-distance migration. Cell 2011;147:1171–85. https://doi.org/10.1016/j.cell.2011.09.052.

37. Wu M, Sun Z, Luo G, Hu C, Zhang W, Han Z. Cloning and characterization of piggyBac-like elements in lepidopteran insects. Genetica 2011;139:149–54. https://doi.org/10.1007/s10709-010-9542-0.

38. Sormacheva I, Smyshlyaev G, Mayorov V, Blinov A, Novikov A, Novikova O. Vertical evolution and horizontal transfer of CR1 non-LTR retrotransposons and Tc1/mariner DNA transposons in Lepidoptera species. Mol Biol Evol 2012;29:3685–702. https://doi.org/10.1093/molbev/mss181.

39. Reiss D, Mialdea G, Miele V, de Vienne D, Peccoud J, Gilbert C, et al. Global survey of mobile DNA horizontal transfer in arthropods reveals Lepidoptera as a prime hotspot. PLOS Genet 2019;15:e1007965. https://doi.org/10.1371/journal.pgen.1007965.

40. Talla V, Suh A, Kalsoom F, Dinc V, Vila R, Friberg M, et al. Rapid increase in genome size as a consequence of transposable element hyperactivity in wood-white (leptidea) butterflies. Genome Biol Evol 2017;9:2491–505. https://doi.org/10.1093/gbe/evx163.

41. Hof AEV t., Campagne P, Rigden DJ, Yung CJ, Lingley J, Quail MA, et al. The industrial melanism mutation in British peppered moths is a transposable element. Nature 2016;534:102–5. https://doi.org/10.1038/nature17951.

42. Gu L, Reilly PF, Lewis JJ, Reed RD, Andolfatto P, Walters JR. Dichotomy of dosage compensation along the neo Z chromosome of the Monarch butterfly. Curr Biol 2019;29:4071–7.

43. Chénais B, Caruso A, Hiard S, Casse N. The impact of transposable elements on eukaryotic genomes: From genome size increase to genetic adaptation to stressful environments. Gene 2012;509:7–15. https://doi.org/10.1016/j.gene.2012.07.042.

44. Osanai-Futahashi M, Suetsugu Y, Mita K, Fujiwara H. Genome-wide screening and characterization of transposable elements and their distribution analysis in the silkworm, Bombyx mori. Insect Biochem Mol Biol 2008;38:1046–57. https://doi.org/10.1016/j.ibmb.2008.05.012.

45. Kapusta A, Suh A, Feschotte C. Dynamics of genome size evolution in birds and mammals. Proc Natl Acad Sci U S A 2017;114:E1460–9. https://doi.org/10.1073/pnas.1616702114.

46. Kidwell MG. Transposable elements and the evolution of genome size in eukaryotes.Genetica 2002;115:49–63. https://doi.org/10.1023/A:1016072014259.

47. Stritt C, Gordon SP, Wicker T, Vogel JP, Roulin AC. Recent activity in expanding populations and purifying selection have shaped transposable element landscapes across natural accessions of the mediterranean grass brachypodium distachyon. Genome Biol Evol 2018;10:304–18. https://doi.org/10.1093/gbe/evx276.

48. Jurka J, Kapitonov V V., Pavlicek A, Klonowski P, Kohany O, Walichiewicz J. Repbase Update, a database of eukaryotic repetitive elements. Cytogenet Genome Res 2005;110:462–7. https://doi.org/10.1159/000084979.

49. Hubley R, Finn RD, Clements J, Eddy SR, Jones TA, Bao W, et al. The Dfam database of repetitive DNA families. Nucleic Acids Res 2016;44:D81–9. https://doi.org/10.1093/nar/gkv1272.

50. Baril T, Imrie R, Hayward A. TobyBaril/EarlGrey: Earl Grey v1.1 2021. https://doi.org/10.5281/ZENODO.5675911.

51. Singh KS, De-Kayne R, Omufwoko KS, Martins DJ, Bass C, ffrench-Constant R, et al. Genome assembly of Danaus chrysippus and comparison with the Monarch Danaus plexippus. G3 Genes|Genomes|Genetics 2021. https://doi.org/10.1093/G3JOURNAL/JKAB449.

52. Wiemers M, Chazot N, Wheat CW, Schweiger O, Wahlberg N. A complete time- calibrated multi-gene phylogeny of the European butterflies. Zookeys 2020;938:97. https://doi.org/10.3897/ZOOKEYS.938.50878.

53. Sultana T, Zamborlini A, Cristofari G, Lesage P. Integration site selection by retroviruses and transposable elements in eukaryotes. Nat Rev Genet 2017;18:292–308. https://doi.org/10.1038/nrg.2017.7.

54. Sawicka K, Bushell M, Spriggs KA, Willis AE. Polypyrimidine-tract-binding protein: A multifunctional RNA-binding protein. Biochem Soc Trans 2008;36:641–7. https://doi.org/10.1042/BST0360641.

55. Tree DRP, Shulman JM, Rousset R, Scott MP, Gubb D, Axelrod JD. Prickle mediates feedback amplification to generate asymmetric planar cell polarity signaling. Cell 2002;109:371–81. https://doi.org/10.1016/S0092-8674(02)00715-8.

56. Laugier E, Yang Z, Fasano L, Kerridge S, Vola C. A critical role of teashirt for patterning the ventral epidermis is masked by ectopic expression of tiptop, a paralog of teashirt in Drosophila. Dev Biol 2005;283:446–58. https://doi.org/10.1016/J.YDBIO.2005.05.005.

57. Datta RR, Lurye JM, Kumar JP. Restriction of ectopic eye formation by Drosophila teashirt and tiptop to the developing antenna. Dev Dyn 2009;238:2202–10. https://doi.org/10.1002/DVDY.21927.

58. Contreras EG, Palominos T, Glavic Á, Brand AH, Sierralta J, Oliva C. The transcription factor SoxD controls neuronal guidance in the Drosophila visual system. Sci Reports 2018 81 2018;8:1–12. https://doi.org/10.1038/s41598-018-31654-5.

59. Matsubara D, Horiuchi SY, Shimono K, Usui T, Uemura T. The seven-pass transmembrane cadherin Flamingo controls dendritic self-avoidance via its binding to a LIM domain protein, Espinas, in Drosophila sensory neurons. Genes Dev 2011;25:1982–96. https://doi.org/10.1101/GAD.16531611.

60. Bessa J, Gebelein B, Pichaud F, Casares F, Mann RS. Combinatorial control of Drosophila eye development by Eyeless, Homothorax, and Teashirt. Genes Dev 2002;16:2415–27. https://doi.org/10.1101/GAD.1009002.

61. Wicker T, Sabot F, Hua-Van A, Bennetzen JL, Capy P, Chalhoub B, et al. A unified classification system for eukaryotic transposable elements. Nat Rev Genet 2007;8:973–82. https://doi.org/10.1038/nrg2165.

62. Oliveira SG, Bao W, Martins C, Jurka J. Horizontal transfers of Mariner transposons between mammals and insects. Mob DNA 2012;3:1–6. https://doi.org/10.1186/1759-8753-3-14.

63. Ivancevic AM, Kortschak RD, Bertozzi T, Adelson DL. Horizontal transfer of BovB and L1 retrotransposons in eukaryotes. Genome Biol 2018;19:1–13. https://doi.org/10.1186/s13059-018-1456-7.

64. Gilbert C, Schaack S, Pace JK, Brindley PJ, Feschotte C. A role for host-parasite interactions in the horizontal transfer of transposons across phyla. Nature 2010;464:1347–50. https://doi.org/10.1038/nature08939.

65. Wallau GL, Ortiz MF, Loreto ELS. Horizontal transposon transfer in eukarya: Detection, bias, and perspectives. Genome Biol Evol 2012;4:689–99. https://doi.org/10.1093/gbe/evs055.

66. Espeland M, Breinholt J, Willmott KR, Warren AD, Vila R, Toussaint EFA, et al. A Comprehensive and Dated Phylogenomic Analysis of Butterflies. Curr Biol 2018;28:770–778.e5. https://doi.org/10.1016/J.CUB.2018.01.061.

67. Ustyugova S V, Amosova AL, Lebedev YB, Sverdlov ED. Cell line fingerprinting using retroelement insertion polymorphism. Biotechniques 2005;38:561–5.

68. Lavoie CA, II RNP, Novick PA, Counterman BA, Ray DA. Transposable element evolution in Heliconius suggests genome diversity within Lepidoptera. Mob DNA 2013;21:1–10.

69. Blass E, Bell M, Boissinot S. Accumulation and rapid decay of non-LTR retrotransposons in the genome of the three-spine stickleback. Genome Biol Evol 2012;4:687–702.

70. Ivancevic AM, Kortschak RD, Bertozzi T, Adelson DL. LINEs between Species: Evolutionary Dynamics of LINE-1 Retrotransposons across the Eukaryotic Tree of Life. Genome Biol Evol 2016;8:3301–22. https://doi.org/10.1093/GBE/EVW243.

71. Evgen’ev MB, Arkhipova IR. Penelope-like elements - a new class of retroelements: distribution, function and possible evolutionary significance. Cytogenet Genome Res 2005;110:510–21. https://doi.org/10.1159/000084984.

72. Kapitonov V V., Tempel S, Jurka J. Simple and fast classification of non-LTR retrotransposons based on phylogeny of their RT domain protein sequences. Gene 2009;448:207. https://doi.org/10.1016/J.GENE.2009.07.019.

73. Woronik A, Tunström K, Perry MW, Neethiraj R, Stefanescu C, Celorio-Mancera M de la P, et al. A transposable element insertion is associated with an alternative life history strategy. Nat Commun 2019 101 2019;10:1–11. https://doi.org/10.1038/s41467-019-13596-2.

74. Hill T, Koseva BS, Unckless RL. The Genome of Drosophila innubila Reveals Lineage-Specific Patterns of Selection in Immune Genes. Mol Biol Evol 2019;36:1405–17. https://doi.org/10.1093/MOLBEV/MSZ059.

75. Sackton TB, Lazzaro BP, Schlenke TA, Evans JD, Hultmark D, Clark AG. Dynamic evolution of the innate immune system in Drosophila. Nat Genet 2007 3912 2007;39:1461–8. https://doi.org/10.1038/ng.2007.60.

76. Shultz AJ, Sackton TB. Immune genes are hotspots of shared positive selection across birds and mammals. Elife 2019;8. https://doi.org/10.7554/ELIFE.41815.

77. Kimbrell DA, Beutler B. The evolution and genetics of innate immunity. Nat Rev Genet 2001 24 2001;2:256–67. https://doi.org/10.1038/35066006.

78. Jurka J, Kapitonov V V., Kohany O, Jurka M V. Repetitive Sequences in Complex Genomes: Structure and Evolution. Annu Rev Genomics Hum Genet 2007;8:241–59. https://doi.org/10.1146/annurev.genom.8.080706.092416.

79. Kidwell MG, Lisch DR. Transposable elements and host genome evolution. Trends Ecol Evol 2000;15:95–9. https://doi.org/10.1016/S0169-5347(99)01817-0.

80. Biémont C, Vieira C. Junk DNA as an evolutionary force. Nature 2006;443:521–4.

81. Cosby RL, Chang N-C, Feschotte C. Host-transposon interactions: conflict, cooperation, and cooption. Genes Dev 2019;33:1098–116. https://doi.org/10.1101/GAD.327312.119.

82. Casacuberta E, González J. The impact of transposable elements in environmental adaptation. Mol Ecol 2013;22:1503–17. https://doi.org/10.1111/mec.12170.

83. Huff JT, Zilberman D, Roy SW. Mechanism for DNA transposons to generate introns on genomic scales. Nature 2016;538:533–6.

84. Gilbert C, Feschotte C. Horizontal acquisition of transposable elements and viral sequences: patterns and consequences. Curr Opin Genet Dev 2018;49:15–24. https://doi.org/10.1016/j.gde.2018.02.007.

85. Galbraith JD, Ludington AJ, Sanders KL, Suh A, Adelson DL. Horizontal transfer and subsequent explosive expansion of a DNA transposon in sea kraits (Laticauda). Biol Lett 2021;17:20210342. https://doi.org/10.1098/RSBL.2021.0342.

86. Le Rouzic A, Capy P. The first steps of transposable elements invasion: Parasitic strategy vs. genetic drift. Genetics 2005;169:1033–43. https://doi.org/10.1534/genetics.104.031211.

87. Kofler R, Senti KA, Nolte V, Tobler R, Schlötterer C. Molecular dissection of a natural transposable element invasion. Genome Res 2018;28:824–35. https://doi.org/10.1101/gr.228627.117.

88. Dupeyron M, Baril T, Hayward A. Broadscale evolutionary analysis of eukaryotic DDE transposons. BioRxiv 2021:2021.09.26.461848. https://doi.org/10.1101/2021.09.26.461848.

89. Prysby MD, Oberhauser KS. Temporal and Geographic Variation in Monarch Densities: Citizen Scientists Document Monarch Population Patterns. Monarch Butterfly Biol Conserv 2004:9.

90. Tang Z, Zhang HH, Huang K, Zhang XG, Han MJ, Zhang Z. Repeated horizontal transfers of four DNA transposons in invertebrates and bats. Mob DNA 2015;6:1–10. https://doi.org/10.1186/s13100-014-0033-1.

91. Kelleher ES, Azevedo RBR, Zheng Y. The evolution of small-RNA-mediated silencing of an invading transposable element. Genome Biol Evol 2018;10:3038–57. https://doi.org/10.1093/gbe/evy218.

92. Goodier JL. Restricting retrotransposons: A review. Mob DNA 2016;7. https://doi.org/10.1186/s13100-016-0070-z.

93. Ray DA, Grimshaw JR, Halsey MK, Korstian JM, Osmanski AB, Sullivan KAM, et al. Simultaneous TE Analysis of 19 Heliconiine Butterflies Yields Novel Insights into Rapid TE-Based Genome Diversification and Multiple SINE Births and Deaths. Genome Biol Evol 2019;11:2162–77. https://doi.org/10.1093/gbe/evz125.

94. Levin HL, Moran J V. Dynamic interactions between transposable elements and their hosts. Nat Rev Genet 2011;12:615–27. https://doi.org/10.1038/nrg3030.

95. Fontanillas P, Hartl DL, Reuter M. Genome organization and gene expression shape the transposable element distribution in the Drosophila melanogaster euchromatin. PLoS Genet 2007;3:2256–67. https://doi.org/10.1371/journal.pgen.0030210.

96. Mawaribuchi S, Takahashi S, Wada M, Uno Y, Matsuda Y, Kondo M, et al. Sex chromosome differentiation and the W- and Z-specific loci in Xenopus laevis. Dev Biol 2017;426:393–400. https://doi.org/10.1016/J.YDBIO.2016.06.015.

97. Dechaud C, Volff JN, Schartl M, Naville M. Sex and the TEs: transposable elements in sexual development and function in animals. Mob DNA 2019 101 2019;10:1–15. https://doi.org/10.1186/S13100-019-0185-0.

98. Erlandsson R, Wilson JF, Päad bo S. Sex Chromosomal Transposable Element Accumulation and Male-Driven Substitutional Evolution in Humans. Mol Biol Evol 2000;17:804–12. https://doi.org/10.1093/OXFORDJOURNALS.MOLBEV.A026359.

99. Chaturvedi P, Hengeveld R, Zechel MA, Lee-Chan E, Singh B. The functional role of class II-associated invariant chain peptide (CLIP) in its ability to variably modulate immune responses. Int Immunol 2000;12:757–65. https://doi.org/10.1093/INTIMM/12.6.757.

100. Minias P, Bateson ZW, Whittingham LA, Johnson JA, Oyler-Mccance S, Dunn PO. Contrasting evolutionary histories of MHC class I and class II loci in grouse-effects of selection and gene conversion. Heredity (Edinb) 2016;116:466. https://doi.org/10.1038/HDY.2016.6.

101. Eizaguirre C, Lenz TL, Kalbe M, Milinski M. Rapid and adaptive evolution of MHC genes under parasite selection in experimental vertebrate populations. Nat Commun 2012 31 2012;3:1–6. https://doi.org/10.1038/ncomms1632.

102. Parfrey LW, Lahr DJG, Katz LA. The dynamic nature of eukaryotic genomes. Mol Biol Evol 2008;25:787–94. https://doi.org/10.1093/molbev/msn032.

103. Sayers EW, Agarwala R, Bolton EE, Brister JR, Canese K, Clark K, et al. Database resources of the National Center for Biotechnology Information. Nucleic Acids Res 2014;42:D7–17. https://doi.org/10.1093/nar/gky1069.

104. Baril T, Imrie R, Hayward A. Earl Grey v1.2 2021. https://doi.org/10.5281/ZENODO.5718734.

105. Smit AFA, Hubley RR, Green PR. RepeatMasker Open-4.0. http://RepeatmaskerOrg 2013.

106. Smit A, Hubley R. RepeatModeler Open-1.0. http://RepeatmaskerOrg 2015.

107. Bao Z, Eddy SR. Automated De Novo Identification of Repeat Sequence Families in Sequenced Genomes. Genome Res 2002;12:1269–76. https://doi.org/10.1101/gr.88502.

108. Price AL, Jones NC, Pevzner PA. De novo identification of repeat families in large genomes. Bioinformatics 2005;21:i351–8.

109. Camacho C, Coulouris G, Avagyan V, Ma N, Papadopoulos J, Bealer K, et al. BLAST+: Architecture and applications. BMC Bioinformatics 2009;10:1-9. https://doi.org/10.1186/1471-2105-10-421.

110. Katoh K, Standley DM. MAFFT multiple sequence alignment software version 7: Improvements in performance and usability. Mol Biol Evol 2013;30:772–80. https://doi.org/10.1093/molbev/mst010.

111. Capella-Gutiérrez S, Silla-Martínez JM, Gabaldón T. trimAl: A tool for automated alignment trimming in large-scale phylogenetic analyses. Bioinformatics 2009;25:1972–3. https://doi.org/10.1093/bioinformatics/btp348.

112. Rice P, Longden L, Bleasby A. EMBOSS: The European Molecular Biology Open Software Suite. Trends Genet 2000;16:276–7. https://doi.org/10.1016/S0168-9525(00)02024-2.

113. Goubert C, Craig RJ, Bilat AF, Peona V, Vogan AA, Protasio A V. A beginner’s guide to manual curation of transposable elements. Mob DNA (In Press 2022:1-98.

114. Larsson A. AliView: a fast and lightweight alignment viewer and editor for large datasets. Bioinformatics 2014;30:3276–8.

115. Fu L, Niu B, Zhu Z, Wu S, Li W. CD-HIT: accelerated for clustering the next- generation sequencing data. Bioinformatics 2012;28:3150–2.

116. Wheeler TJ, Eddy SR. nhmmer: DNA homology search with profile HMMs. Bioinformatics 2013;29:2487–9. https://doi.org/10.1093/BIOINFORMATICS/BTT403.

117. Xu Z, Wang H. LTR_FINDER: an efficient tool for the prediction of full-length LTR retrotransposons. Nucleic Acids Res 2007;35:W265–8. https://doi.org/10.1093/nar/gkm286.

118. Ou S, Jiang N. LTR_FINDER_parallel: parallelization of LTR_FINDER enabling rapid identification of long terminal repeat retrotransposons. BioRxiv 2019:2–6.

119. Wong WY, Simakov O. RepeatCraft: a meta-pipeline for repetitive element de- fragmentation and annotation. Bioinformatics 2018;35:1051–2. https://doi.org/10.1093/bioinformatics/bty745.

120. Rubino F, Creevey CJ. MGkit: Metagenomic framework for the study of microbial communities 2014.

121. Edgar RC. MUSCLE: Multiple sequence alignment with high accuracy and high throughput. Nucleic Acids Res 2004;32:1792–7. https://doi.org/10.1093/nar/gkh340.

122. Lenhard B, Sandelin A, Carninci P. Metazoan promoters: Emerging characteristics and insights into transcriptional regulation. Nat Rev Genet 2012;13:233–45. https://doi.org/10.1038/nrg3163.

123. Quinlan AR, Hall IM. BEDTools: A flexible suite of utilities for comparing genomic features. Bioinformatics 2010;26:841–2. https://doi.org/10.1093/bioinformatics/btq033.

124. Makowski D, Ben-Shachar M, Lüdecke D. bayestestR: Describing Effects and their Uncertainty, Existence and Significance within the Bayesian Framework. J Open Source Softw 2019;4:1541. https://doi.org/10.21105/joss.01541.

125. Gel B, Serra E. karyoploteR: an R/Bioconductor package to plot customizable genomes displaying arbitrary data. Bioinformatics 2017;33:3088–90. https://doi.org/10.1093/bioinformatics/btx346.

126. Kimura M. A simple method for estimating evolutionary rates of base substitutions through comparative studies of nucleotide sequences. J Mol Evol 1980;16:111–20.

127. Pfeiler E, Nazario-Yepiz NO, Pérez-Gálvez F, Chávez-Mora CA, Laclette MRL, Rendón-Salinas E, et al. Population Genetics of Overwintering Monarch Butterflies, Danaus plexippus (Linnaeus), from Central Mexico Inferred from Mitochondrial DNA and Microsatellite Markers. J Hered 2017;108:163–75. https://doi.org/10.1093/jhered/esw071.

128. Wickham H. ggplot2: elegant graphics for data analysis. Springer; 2016.

129. Wickham H, Averick M, Bryan J, Chang W, McGowan LD, François R, et al. Welcome to the Tidyverse. J Open Source Softw 2019;4:1686.

130. Marchler-Bauer A, Bo Y, Han L, He J, Lanczycki CJ, Lu S, et al. CDD/SPARCLE: functional classification of proteins via subfamily domain architectures. Nucleic Acids Res 2017;45:D200–3. https://doi.org/10.1093/NAR/GKW1129.

131. Evgen’ev MB, Zelentsova H, Shostak N, Kozitsina M, Barskyi V, Lankenau DH, et al. Penelope, a new family of transposable elements and its possible role in hybrid dysgenesis in Drosophila virilis. Proc Natl Acad Sci 1997;94:196–201. https://doi.org/10.1073/PNAS.94.1.196.

132. Nguyen L-T, Schmidt HA, von Haeseler A, Minh BQ. IQ-TREE: A Fast and Effective Stochastic Algorithm for Estimating Maximum-Likelihood Phylogenies. Mol Biol Evol 2015;32:268–74. https://doi.org/10.1093/MOLBEV/MSU300.

133. Kalyaanamoorthy S, Minh BQ, Wong TKF, Haeseler A von, Jermiin LS. ModelFinder: fast model selection for accurate phylogenetic estimates. Nat Methods 2017 146 2017;14:587–9. https://doi.org/10.1038/nmeth.4285.

134. Team RC. R: A language and environment for statistical computing 2013.

