## Additional File 2 for "Migrators Within Migrators: Exploring Transposable Element Dynamics in the Monarch Butterfly, *Danaus plexippus*"

A recent study focussing on a new genome assembly of the African queen butterfly, *Danaus chrysippus,* found repeat content in the genome assembly for the monarch butterfly, *Danaus plexippus*, to be between 11.2% and 14.3% [1]. In contrast, we found repeat content in the same assembly to be 6.21%, approximately half that identified in [1]. Consequently, we investigated the discrepancies between the results of [1] and those presented in our study. We provide a detailed summary below, so that disparities in methodology underlying the large difference in reported repeat content are fully transparent, allowing repeatability and traceability.

Firstly, the final masking of the monarch assembly in our study was performed with a conservative score threshold of 400 to exclude poor matches unlikely to be true TE sequences (see methods). A final step also involved removing annotations <100bp to decrease the incidence of spurious hits (see methods). When the same filtering requirements are imposed on the TE annotations of [1], we see a reduction in the total number of annotations from 177,493 elements (30.29Mb) to 81,537 (22.49Mb), a decrease of 95,956 annotations (7.8Mb, or 46% or all annotations). Thus, almost half the annotations in [1] are low scoring and very short sequences. We also filter repeat annotations to remove overlapping TE annotations, as a single base pair cannot be attributed to multiple repeat classifications. When removing overlaps from the filtered annotations from [1], annotations are further reduced from 81,537 (22.49Mb) to 69,281 (19.51Mb), resulting in a total loss of 108,212 annotations (10.78Mb). These steps alone reduce the repeat annotation from 12.18% to 7.85% of the monarch genome. Our methodology also uses defragmentation of repeat annotations using RepeatCraft (see methods). Consequently, the mean length of TE annotation in our results is 353bp, whilst the results of [1] are more fragmented, with a mean length of 223bp.

In [1], a *de novo* TE library was generated by analysing the assembly of *D. chrysippus* using a basic RepeatModeler run with no downstream quality filtering, clustering, or sequence validation [1]. Following this, the *de novo* library was combined with an undefined Lepidoptera library from RepeatMasker v4.1.0 (it is unknown if this is the RepBase or Dfam library, or a combination, and which versions) and this was used to annotate the monarch genome assembly [1]. The TE libraries used in our study were generated using the genome assembly of interest, rather than applying a library generated from a single genome (i.e. *D. chrysippus* in [1]) to annotate multiple species. Thus, for the monarch, we used the monarch genome assembly to detect novel repeats, which we believe is an important methodological requirement for generating accurate TE annotations.

Following automated *de novo* curation, putative TE sequences were manually curated according to established manual curation protocols (see methods). Manual curation enables novel TE classification based on structural, sequence, and protein domain evidence to classify TEs with higher confidence than automated tools. When investigating the TE library for *D. chrysippus* in [1], we found a high level of redundancy, with 2,906 sequences grouping into 2,103 clusters with CD-Hit-est, using the library clustering parameters specified in a recent paper on manual curation (i.e. -d 0 -aS 0.8 -c 0.8 -G 0 -g 1 -b 500) [2]. This redundancy has the potential to inflate copy number estimates by annotating separate fragments of the same TE with different families, which is avoided in our study by removing library redundancy through clustering followed by defragmentation of repeat annotations to identify fragments that are likely to be from the same initial insertion. Furthermore, 25% (403) of the *de novo* library sequences in [1] are very short (<300bp), with 85 being <100bp. Of these, 70 are classified as LINEs, although due to their short length it is unlikely they can be classified with high confidence.

Considering high copy number TE families that were not identified in our study, the highest copy number family in [1] was classified as a rolling circle family. However, searches against CDD [3] and RepBase using the CENSOR web tool (https://www.girinst.org/censor/) [4,5] revealed no evidence to classify the family as such, and annotations in the genome were very short, with a mean length of 141bp. The next three highest copy number families in [1] also returned no hits in CDD or CENSOR, although these families were classified as a LINE family, an unclassified element family, and a rolling circle family, respectively. Furthermore, the longest single annotation reported in [1] that was missing from our TE library had 16 short fragmented hits to TEs of varying classifications in CENSOR, including DNA, rolling circle, LTR, and LINE hits in stretches ~40-80bp across the consensus sequence, although it was labelled in the library as a LINE (Figure 1).

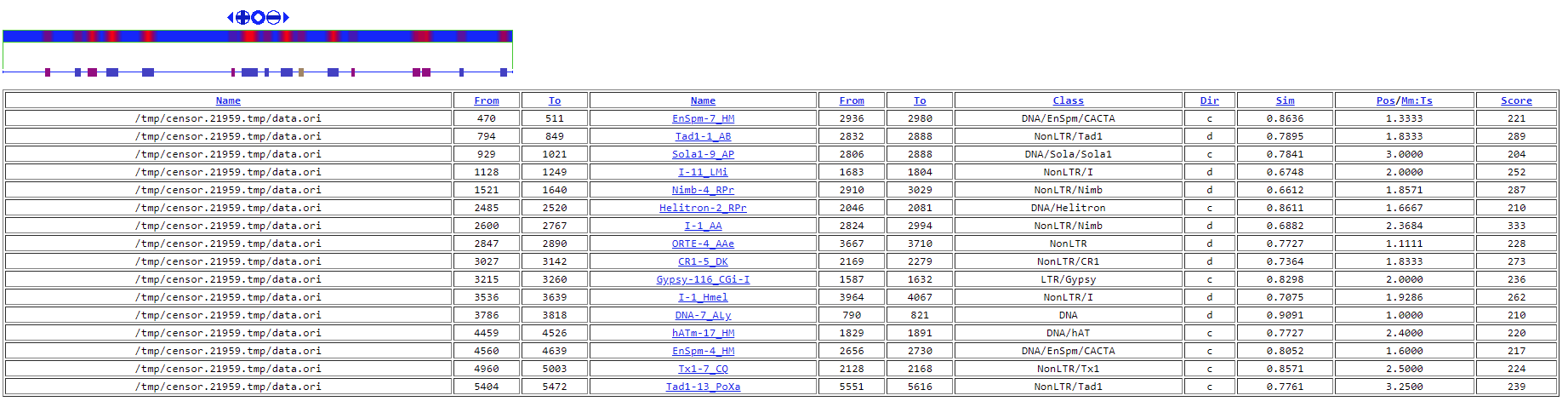

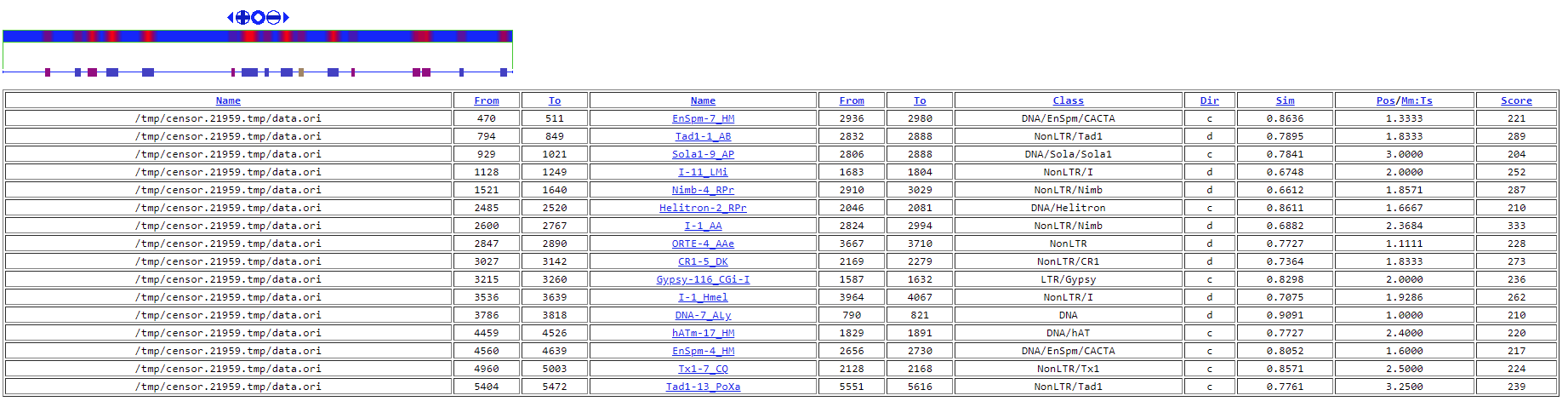

Figure 1. CENSOR hits to a *de novo* sequence classified as a LINE in [1]. The top karyoplot shows hits to known TEs as red bars, while the blue bar is consensus sequence. The table shows hits to known TEs across the consensus sequence, including hits to DNA, NonLTR, Helitron, and LTR elements.

In summary, the greater reported repeat content in [1] appears to arise due to quality filtering and classification validation issues, in combination with the use of lower scoring annotation thresholds, and the inclusion of overlapping and highly fragmented annotations. Collectively these issues are likely to have resulted in overestimated TE copy number and TE coverage for the monarch genome in [1].

References

[1]	Singh KS, De-Kayne R, Omufwoko KS, Martins DJ, Bass C, ffrench-Constant R, et al. Genome assembly of Danaus chrysippus and comparison with the Monarch Danaus plexippus. G3 Genes|Genomes|Genetics 2021. https://doi.org/10.1093/G3JOURNAL/JKAB449.

[2]	Goubert C, Craig RJ, Bilat AF, Peona V, Vogan AA, Protasio A V. A beginner’s guide to manual curation of transposable elements. Mob DNA (In Press 2022:1–98.

[3]	Marchler-Bauer A, Bo Y, Han L, He J, Lanczycki CJ, Lu S, et al. CDD/SPARCLE: functional classification of proteins via subfamily domain architectures. Nucleic Acids Res 2017;45:D200–3. https://doi.org/10.1093/NAR/GKW1129.

[4]	Jurka J, Klonowski P, Dagman V, Pelton P. Censor—a program for identification and elimination of repetitive elements from DNA sequences. Comput Chem 1996;20:119–21. https://doi.org/10.1016/S0097-8485(96)80013-1.

[5]	Kohany O, Gentles AJ, Hankus L, Jurka J. Annotation, submission and screening of repetitive elements in Repbase: RepbaseSubmitter and Censor. BMC Bioinformatics 2006;7:1–7. https://doi.org/10.1186/1471-2105-7-474/FIGURES/2.
