## Supplementary figures and images for "Migrators Within Migrators: Exploring Transposable Element Dynamics in the Monarch Butterfly, *Danaus plexippus*"

### Additional File 3

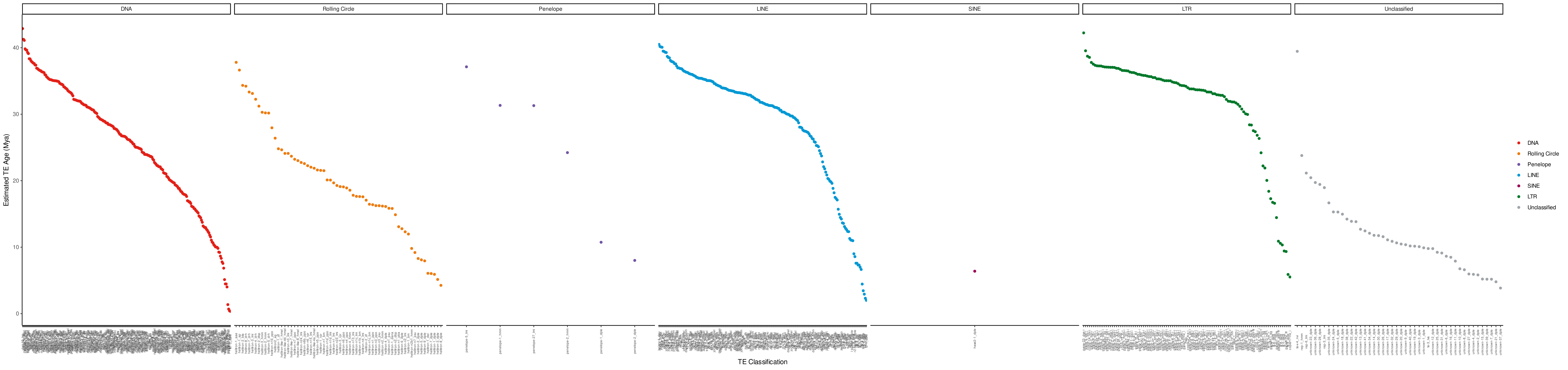

### Additional File 4

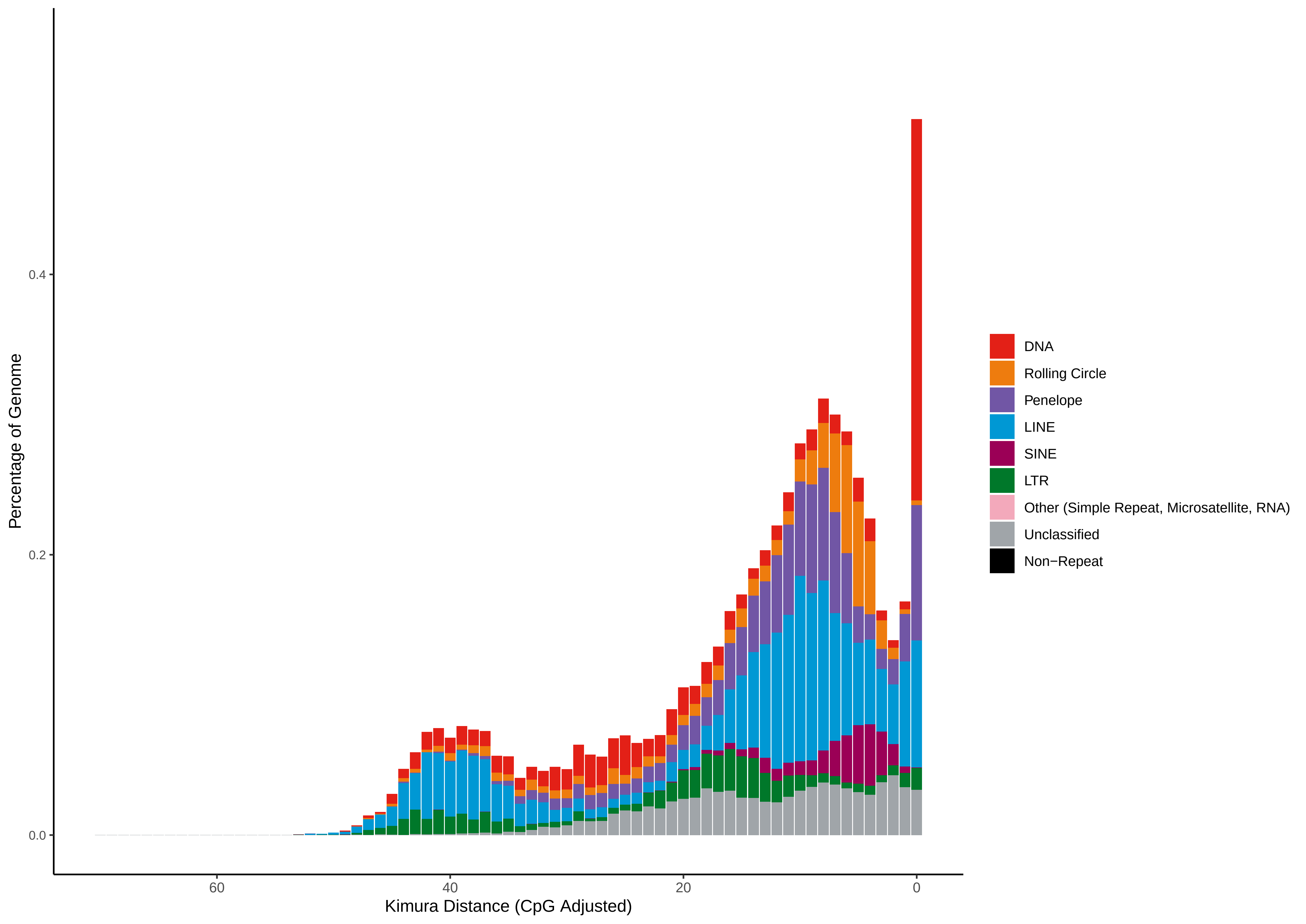

### Additional File 5

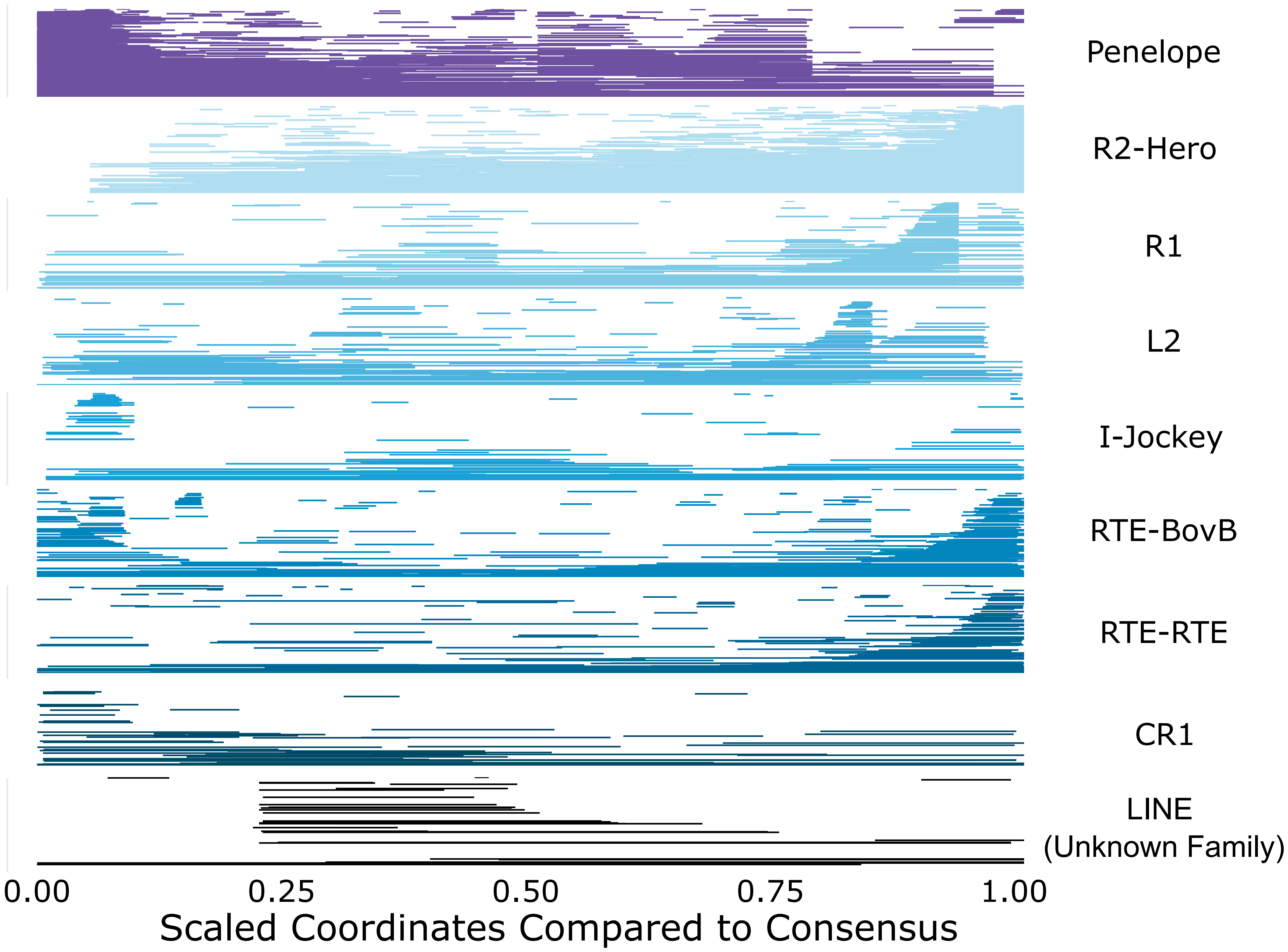

### Additional File 6

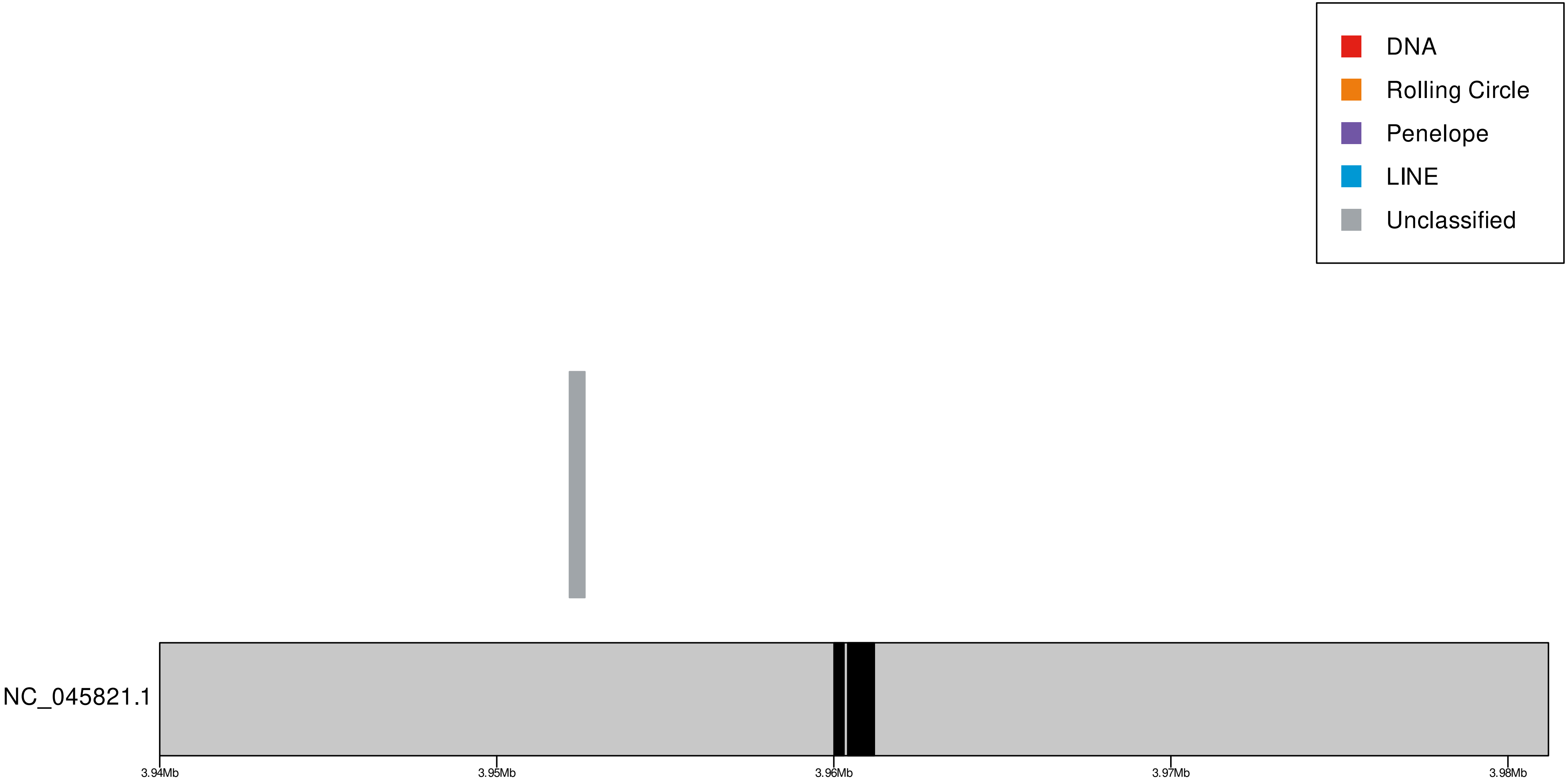

### Additional File 7

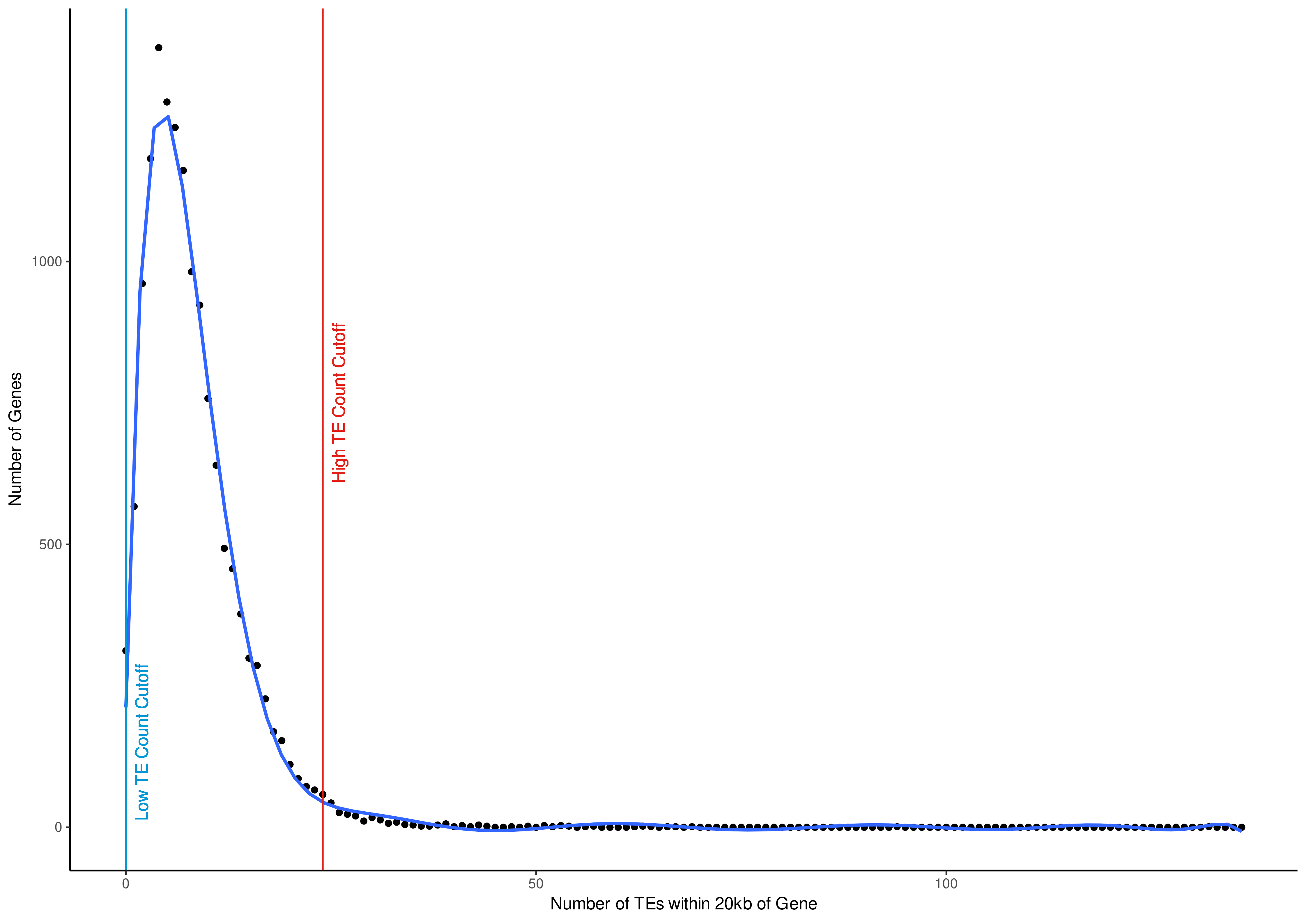

### Additional File 8

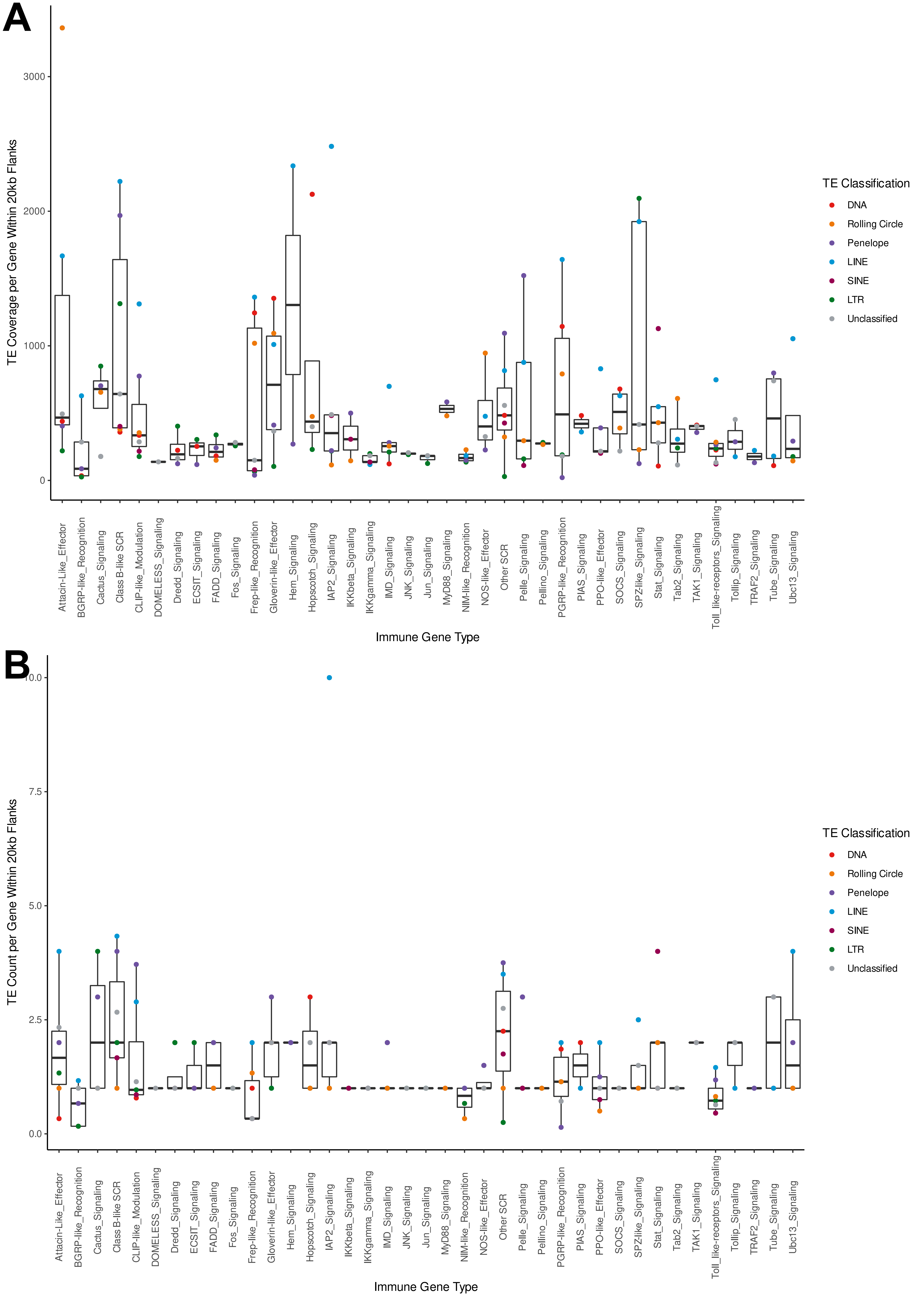

### Additional File 9

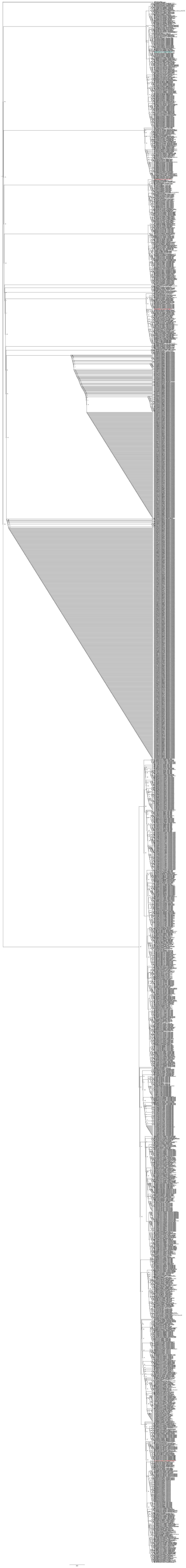

### Additional File 13

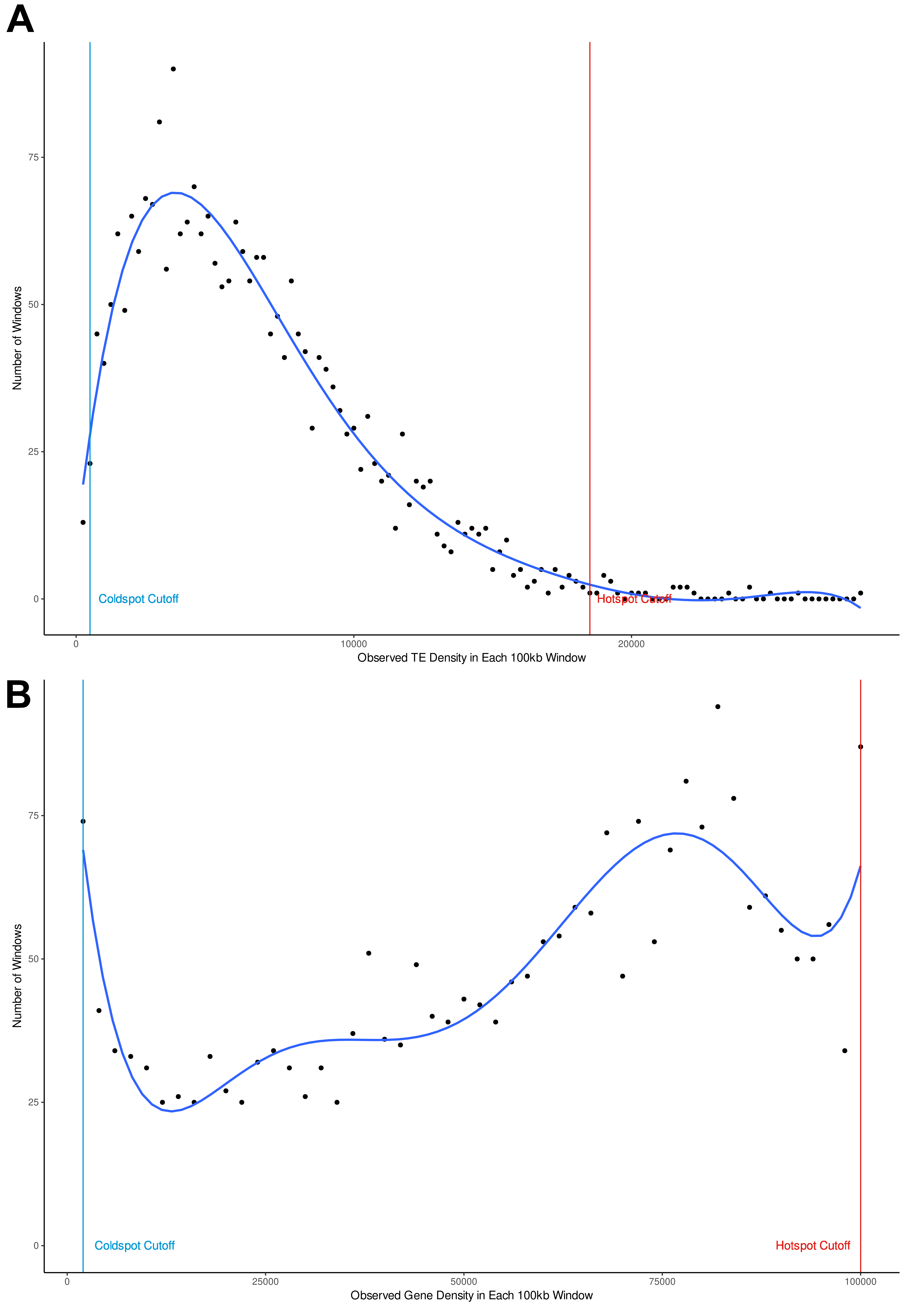
